# Investigating unexplained genetic variation and its expression in the arbuscular mycorrhizal fungus *Rhizophagus irregularis*

**DOI:** 10.1101/682385

**Authors:** Frédéric G. Masclaux, Tania Wyss, Marco Pagni, Pawel Rosikiewicz, Ian R. Sanders

## Abstract

Arbuscular mycorrhizal fungi (AMF) are important symbionts of plants. Recently, studies of the AMF *Rhizophagus irregularis* recorded within-isolate genetic variation that does not completely match the proposed homokaryon or heterokaryon state (where heterokaryons comprise a population of two distinct nucleus genotypes). We re-analysed published data showing that bi-allelic sites (and their frequencies), detected in proposed homo- and heterokaryote *R. irregularis* isolates, were similar across independent studies using different techniques. This indicated that observed within-fungus genetic variation was not an artefact of sequencing and that such within-fungus genetic variation possibly exists. We looked to see if bi-allelic transcripts from three *R. irregularis* isolates matched those observed in the genome as this would give a strong indication of whether bi-allelic sites recorded in the genome were reliable variants. In putative homokaryon isolates, very few bi-allelic transcripts matched those in the genome. In a putative heterokaryon, a large number of bi-allelic transcripts matched those in the genome. Bi-allelic transcripts also occurred in the same frequency in the putative heterokaryon as predicted from allele frequency in the genome. Our results indicate that while within-fungus genome variation in putative homokaryon and heterokaryon AMF was highly similar in 2 independent studies, there was little support that this variation is transcribed in homokaryons. In contrast, within-fungus variation thought to be segregated among two nucleus genotypes in a heterokaryon isolate was indeed transcribed in a way that is proportional to that seen in the genome.

## Introduction

Arbuscular mycorrhizal fungi (AMF) colonize the roots of the majority of land plants, improving plant nutrient and water uptake in exchange for plant-assimilated carbohydrates and lipids [1, 2]. Arbuscular mycorrhizal fungi form hyphae and spores that contain a continuous cytoplasm with many co-existing nuclei. There is no known stage during the AMF life cycle in which only one nucleus initiates a new generation, suggesting that a multinucleate organization is important [3].

Over the last two decades there has been some debate about whether AMF are homokaryons (possessing genetically identical nuclei) or heterokaryons (possessing two or more genetically different nuclei). The ensuing, and lengthy, debate about whether or not AMF are heterokaryons has largely confounded two separate questions. 1. Are AMF heterokaryons or homokaryons, without considering the quantity of genetic differences among nuclei? 2. How much genetic variation exists within an AMF isolate that could potentially be distributed among nuclei? Resolving this issue of genetic variation in this fungus is biologically highly relevant because clonally produced single spore offspring of *R. irregularis* show high variation in their phenotype [4] and extremely high variation in their effects on plant growth [5].

Studies over the last two decades that have tried to address the two questions outlined above are highly inconsistent in their conclusions regarding the amount of genetic variation existing within an AMF isolate and the prediction of the number of genetically different nuclei (summarized in Figure 1). However, there has been a lack of studies focused on the same species or isolate. Some species studied probably diverged from each other hundreds of millions of years ago, making comparisons between some of these studies as largely meaningless [6]. Furthermore, different approaches have been used as technologies have developed over the years.

**Figure 1.**
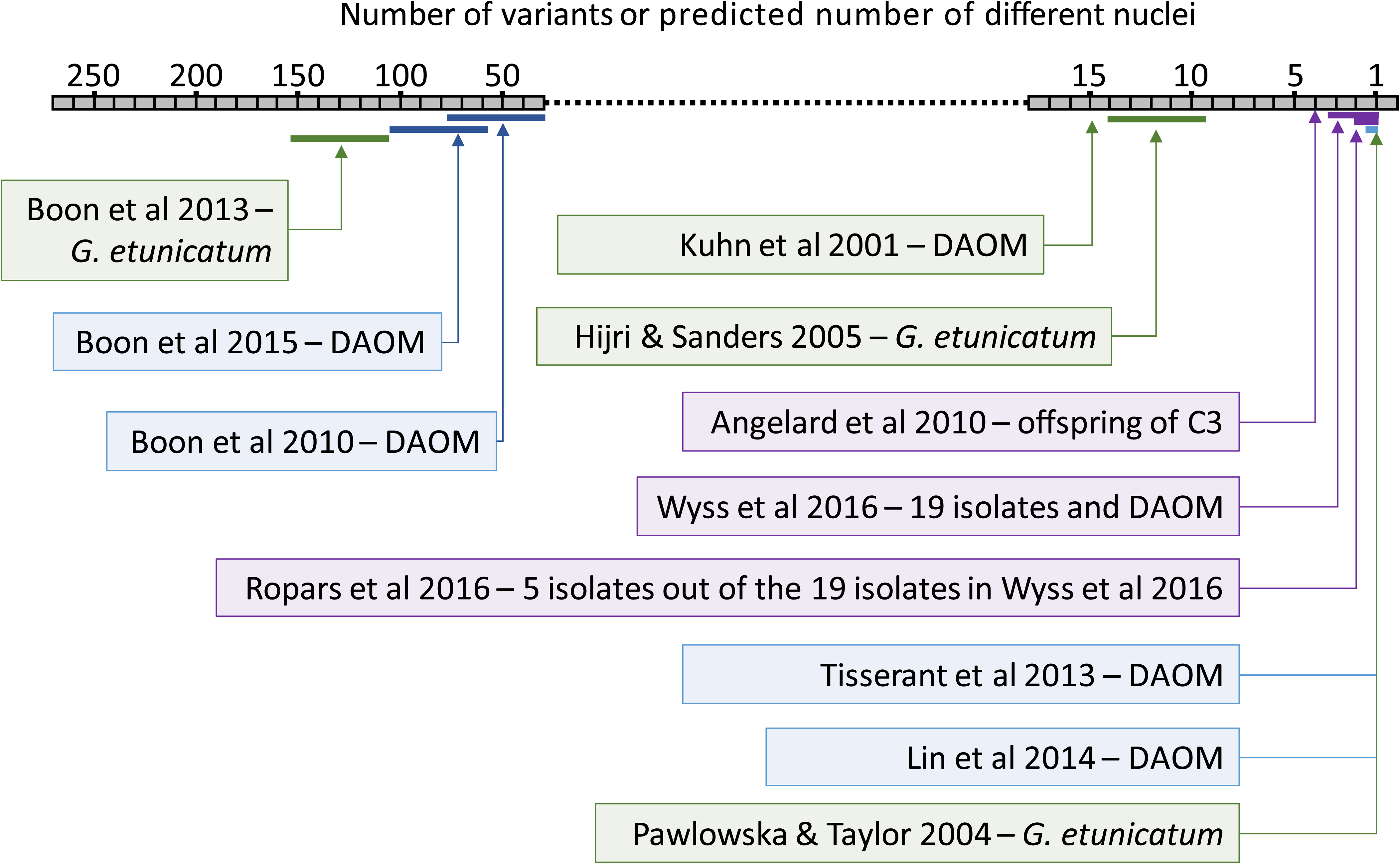
Range of reported levels of intra-isolate genetic variation in different AMF species and different studies. The level of intra-isolate genetic variation is expressed as the number of alleles observed per position in the genome or the predicted number of genetically different nuclei per AMF isolate on the grey horizontal bar. Levels of intra-isolate genetic variation were determined by sequencing methods. Green represents *Glomus etunicatum* although the studies did not use the same isolates. Blue represents *Rhizophagus irregularis* isolate DAOM 197198. Purple represents studies including isolates of *R. irregularis* other than DAOM 197198. It is important to note that within some studies a greater number of alleles was observed compared to the number of different nuclei predicted, owing to assumptions about ploidy or gene copy number.

More recently, studies using different techniques and sequencing platforms have concentrated on describing the genome of one haploid species of AMF, *R. irregularis*, and its within and among isolate variation [7–11]. Thus, there are now directly comparable data originating from independent studies. The whole genome of the *R. irregularis* isolate, DAOM197198, was sequenced showing that within fungus polymorphism existed but was low. The authors concluded that this isolate was a homokaryon [7, 12] and did not try and explain the variation. Double-digest restriction site-associated sequencing (ddRAD-seq) was performed on DAOM197198 and 19 other *R. irregularis* isolates [8], showing different levels of within-fungus polymorphism among the isolates that suggested isolates with a predominantly homokaryon or heterokaryon state; where heterokaryons comprised predominantly only two different genotypes of nuclei, but with the possibility of additional genetically different nuclei at low frequency. Ropars *et al.* (2016) sequenced the whole genome of five of the 19 isolates and concluded that three of these isolates; A1, B3 and C2 were homokaryons and that two isolates; A4 and A5 were heterokaryons with two genetically different nucleus genotypes [9]. Notably, none of these studies have recorded high levels of polymorphism as those observed by Boon et al. [13, 14] (Figure 1). Despite this, Bruns et al. (2018) [15] suggested a strong dichotomy between, on the one hand, the studies of Boon et al. and Wyss et al. [8, 13, 14] and on the other hand, the studies by Ropars et al., Lin et al. and Tisserant et al. [7, 9, 12] (see discussion by Sanders 2018).

All studies revealed a level of polymorphism unexplainable by a strict homokaryon state. In each case, bi-allelic sites existed in isolates that had been predicted to be homokaryons. However, this variation was largely discounted as being due to sequencing errors. Plots of allele frequency at bi-allelic sites in proposed homokaryons and heterokaryon isolates revealed a peak at 0.5 in several isolates but with additional variation at other frequencies[8]. A clear 0.5 peak of frequency at bi-allelic sites is expected for a heterolaryon with equal proportions of two nucleus genotypes or a diploid. Ropars et al. (2016) found a peak at 0.5 frequency in A4 and A5, but a U-shaped distribution in the other isolates, which is difficult to explain and not consistent with [8, 9].

Wyss et al. (2016) observed within-fungus genetic variation among biological replicates of the same isolates, thereby, reducing the probability that the variation was the result of sequencing error [8]. Ropars et al. (2016) also constructed replicate libraries for whole genome sequencing, although they pooled them for further analysis, and did not use them as replicates[9]. We hypothesized that bi-allelic sites observed in the genome of a given isolate in a replicated study, and between the two independent studies, will be consistent if they did not arise from sequencing error. Thus, observed variation represents true variation and is not due to sequencing error. Using the replicates of Wyss et al. and the data from Ropars et al., treated as replicates, allows to test whether the unexplained variation in these studies is indeed real biological variation or an artefact of sequencing. This was the first goal of this study.

So far there has been almost no attention paid to the functional significance of any within-fungus genome variation observed in AMF. However, single spore offspring of clonally produced *R. irregularis* shows a high degree of variation in their phenotype and their effects on plant growth [4, 5]. Transcriptome profiling of *R. irregularis* would reveal whether the genome variation observed at bi-allelic sites is transcribed or whether unequal or single-allele expression occurs. However, as yet, no studies exist in AMF relating variation at the transcriptome level back to variation observed at the genome level. The detection of bi-allelic transcripts that match observed variation in the genome would confirm the existence of bi-alleles in the genome and would indicate that the variation could potentially have consequences on the phenotype. Expression of two alleles of a given gene, unequal expression of the two alleles, or expression of only one of them, could all have consequences on the phenotype of the fungus or its effect on plant growth. If alleles were truly located on two genetically different nuclei, it would mean that it would also represent a measure of the contribution by the two different AMF nucleus genotypes to overall gene expression in the fungus. In plants, the contribution of two different genomes (for example in hybrids or polyploids) is often highly unequal and the contribution from each genome can be tissue specific [16].

Thus, the existence of two different genotypes of nuclei does not necessarily mean that gene expression will be in equal proportions from the two alleles at a given locus. Thus, the second goal of our study was, therefore, to investigate whether variation observed in the *R. irregularis* transcriptome matched variation observed in the genome as an additional test of whether the variation really exists. Additionally, we wanted to know the contribution of each nucleus genotype to overall gene expression in this fungus.

We, therefore, used the available genome data on *R. irregularis* to compare within-isolate genome polymorphism and test whether the unexplained variation in these studies is indeed real biological variation or an artefact of sequencing. We also conducted transcriptome profiling on the isolate C3 (that has been predicted to be a heterokaryon with two nucleus genotypes) and the isolate C2 (that has been predicted to be a homokaryon) to see if variation found at the genome level is mirrored at the transcriptome level, i.e. whether this variation is co-expressed, and in what proportion the alleles are expressed. Finally, we looked at the extent of structural variation occurring in the genomes of these isolates because structural variation can also strongly affect the phenotype of an organism. Although heterokaryon can refer to the co.-existence of two or more genetically distinct nuclei, here we refer to heterokaryon isolates as those possessing primarily two genetically different nuclei.

## Materials and Methods

### Fungal material, RNA extraction and library preparation

Three *R. irregularis* isolates (C2, C3 and DAOM197198) were cultured *in vitro* with *Ri* T-DNA transformed carrot roots [17] and a split plate system [18]. Fungal material was extracted from the medium in citrate buffer (450 mL ddH_2_O, 8.5 mL 0.1 N citric acid, 41.5 mL 0.1 N Na citrate) for 20 min and washed with sterile double-deionized water (ddH_2_O). Material from three Petri plates was pooled per sample. There were three samples per isolate. RNA was extracted with the RNeasy Plant Mini kit (Qiagen). Two micrograms of RNA from each sample was used to prepare libraries for RNA sequencing (RNA-seq) using the TruSeq Stranded Total RNA Library Prep Kit (Illumina) with a PCR enrichment step of 15 cycles. The libraries were pooled and sequenced using Illumina HiSeq 2500 (100 bp paired-end reads). Sequences were deposited in the NCBI SRA database (BioProject Accession Number: <u>PRJNA509102</u>).

### All data used in this study

The origin and characteristics of whole genome (WG), ddRAD-seq and RNA-seq data from *R. irregularis* and other species that was used in this study is summarized in Tables S1-S3. The name of the isolate “DAOM197198” is shortened to “DAOM” in some of the figures.

Additionally, in order to find examples to test the RNA-seq KisSplice pipeline (described below), we screened the NCBI sequence archive to select RNA-seq data of species that have homokaryon and heterokaryon states or have homozygote and heterozygote states (Table S3). The idea was to find species where both states exist like *Arabidopsis* homozygote (*Col*-0 accession) and *Arabidopsis* heterozygote (Hybrid line). Some other fungal species with known homokaryon and heterokaryon states were also included (Table S3).

### Processing sequence data

Raw reads were processed with the script Tagcleaner.pl to trim Illumina adapters [19]. Reads were processed with PrinSeq-lite.pl version 0.20.4 [20] to remove reads containing ‘Ns’, to trim low quality 3’-ends and retain reads longer than 50 bp. A summary of the read size after filtering is in Table S4.

### In silico prediction of ddRAD-seq fragments

To compare the same sites in ddRAD-seq and WG data, analysis was restricted to ddRAD-seq fragments existing in WG data. In *silico* digestion with *EcoR*I and MseI was performed on each genome assembly to define predicted ddRAD-seq fragments longer than 49 bp and delimited by a *EcoR*I site and a MseI site. Predicted fragments were aligned to their respective genome with Novoalign v3.02.00 (Novocraft-Technologies, 2014). Fragments that could not be re-aligned were removed. Re-aligned predicted fragments define regions that were considered in subsequent analyses. We refer to ‘predicted ddRAD-seq fragment’ as one of these fragments and to ‘predicted ddRAD-seq region’ to designate the location where the fragment mapped (Figure S1).

### Identification of interspersed repeats and repeated elements

We use the term ‘repeats’ to include the interspersed repeats (mobile elements and transposon-like regions) and repeated elements (multicopy genes and duplicated regions). The interspersed repeat families were predicted *de novo* with the program RepeatModeler Open-1.0.8 (http://www.repeatmasker.org) and interspersed repeats were annotated with RepeatMasker Open-4.0.6 (http://www.repeatmasker.org).

Every method of repeat detection has its own stringency, which has direct consequences on the number of bi-allelic positions that can be discovered. We wanted to test that identification of poly-allelic sites was not strongly biased by the method of repeat detection. Thus, we used three methods to identify repeated elements to test the influence of underestimating or overestimating the repeated elements on within-isolate polymorphism.

Method M03 was used previously [8]. The *in silico* predicted ddRAD-seq fragments were submitted to pairwise comparisons using ggsearch36; a global pairwise alignment algorithm from the package fasta-36.3.5e [21]. This was done to identify globally similar predicted fragments among the ddRAD-seq fragments. Fragments having a match with another fragment were labelled ‘repeated elements’. The M03 repeats corresponded to interspersed repeats and M03 repeated elements. Method M03 is the default method used in this study (Figure S1).

Method M07 had a larger spectrum than M03. *In silico* predicted ddRAD-seq fragments were aligned to the whole reference genome using glsearch36 (package fasta-36.3.5e) to identify fragments matching more than one time in the reference genome. Fragments with more than one match were labelled ‘repeated elements’. The M07 repeats corresponded to a combination of interspersed repeats and M07 repeated elements. It is the most stringent of the three methods.

With a third method, M12, *in silico* predicted ddRAD-seq fragments were mapped to the genome assembly with Novoalign V3.02.00 using parameters -r All 100 -R 400 -s0. Fragments with more than one match in the assembly were labelled ‘repeated elements’. The M12 repeats were a combination of interspersed repeats and M12 repeated elements. This method is of medium stringency.

### Gene prediction in genome assemblies

Coding regions were defined with the *ab initio* gene predictor Augustus version 3.1 using the gene model previously defined for the N6 assembly, constructed for the DAOM197198 isolate [8].

### Mapping of WG and ddRAD-seq sequence data on reference genomes

Wyss et al. (2016) mapped reads to the DAOM197198 genome assembly because other genome assemblies of *R. irregularis* isolates were not available at the time [8]. For this study, we used genome assemblies of isolates A1, A4, A5, B3 and C2 for mapping. We aligned reads against the genome assemblies with Novoalign v3.02.12 (Novocraft-Technologies, 2014). The correctly mapped paired-end reads were selected for subsequent analyses using Samtools version 1.3 [22]. Single nucleotide polymorphisms (SNPs), insertions and deletions (indels) and multiple nucleotide polymorphisms (MNPs) were called for each sample using Freebayes version 1.0.2 [23] at positions with a minimum 10× depth of coverage. Only alleles with frequencies greater or equal to 0.1 were recorded. Variant files were filtered to keep the positions with a phred-scaled quality score greater or equal to 30. For every sample, and at each site, we reported whether there was a single allele, two alleles or more than two alleles. A missing value (NA) was assigned if the depth of coverage was below 10×. Because the objective of the study was to study all the within-isolate polymorphism, no filter for the allelic ratio was applied to a position to call it bi-allelic or poly-allelic position.

We compared variants in ddRADseq data [8] and WG data [9]. If the same variants are found among replicates and between two independent studies, then it is highly unlikely that the observed variants are artefacts caused by error generated in the sequencing. Wyss et al. (2016) included 3 replicates per isolate. Ropars et al. (2016) assembled genomes of *R. irregularis* that were pooled samples of different libraries that can also be treated as replicates. The numbers of replicates in WG sequencing was between 2 and 3, depending on the isolate.

### Bi-allelic site density and allele frequency

In order to only consider variants in single copy genes, the variant file of each sample was filtered to only retain positions with two alleles in non-repeated and coding regions. The number of bi-allelic positions was divided by the number of positions from predicted ddRAD-seq regions that were sequenced with a coverage greater or equal to 10.

All positions from the variant file were included except those in repeated regions defined by RepeatModeler/RepeatMasker and with less than 10× coverage. The threshold was altered to test the effect of depth of coverage on allele frequency distribution. The list of all allele frequencies was plotted using the density function from the R package ggplot2.

### Depth of coverage and predicted ddRAD-seq regions

Low coverage in samples would limit the ability to reliably detect polymorphisms. Samtools version 1.3 [22] was used to calculate depth of coverage in ddRAD-seq and WG data. Samtools and custom perl script were used to calculate the mean depth of coverage per predicted ddRAD-seq region. A mean coverage greater, or equal to, 10 was necessary to consider a predicted ddRAD-seq region as covered. A similar method was used to calculate the depth of coverage per position required to calculate bi-allelic position density.

### Identification of common bi-allelic positions

The first goal of this study was to compare whether the same bi-allelic sites were found among replicates and studies. The R package UpSetR [24] was applied on lists of bi-allelic positions in non-repeated, coding regions to find common bi-allelic positions in different replicates and generate UpSet plots.

### Discovery of bi-allelic SNPs in RNA-seq data

KisSplice performs a local assembly of RNA-seq reads and detects variants directly from a De Bruijn graph, independently of a reference genome [25]. The methodology to analyse RNA-seq data is summarized in Figure S2. Because KisSplice reports a very local context around SNPs, we generated reference full-length transcriptome assemblies from RNA-seq data with Trinity [26]. Tritnity v2.3.2 was run with standard parameters (except --SS_lib_type RF). Predicted ORFs were identified in the Trinity assembled transcripts with TransDecoder (v5.0.2). Next, we identified full-length, or nearly full-length, transcripts (https://github.com/trinityrnaseq/trinityrnaseq/wiki/Counting-Full-Length-Trinity-Transcripts, accessed on July 2018). The assembled transcripts were compared to known proteins in Swissprot UniProtKB proteins (www.uniprot.org, accessed on June 2017). The comparison was performed with BLASTX (2.6.0+) with the following parameters: −evalue 1e-20 −max_target_seqs 1 −outfmt 6. The length of the top hit and percentage of the hit length, included in the alignment to the Trinity transcript, were added to the BLASTX result file with the provided script analyze_blastPlus_topHit_coverage.pl. Next, a custom Perl script was applied on the resulting file to keep only the best isoform of each gene and to remove genes with multiple groups (following Trinity’s specifications for ‘group’ and ‘isoform’). KisSplice version 2.4.0 was run on RNA-seq data of each organism with the parameters: --experimental −s 2 −k 41. We focused on the ‘Type0a bubbles’ (KisSplice nomenclature), that are two identical short sequences differing by a single SNP at a given position. Bubble sequences were positioned on the Trinity-reference transcriptome using BLAT with the option -minIdentity=80. To predict amino acid changes of a SNP, the program KisSplice2RefTranscriptome (K2RT) was run taking as input the predicted ORFs, the Type0a bubbles produced by KisSplice, and the mapping results of BLAT. A custom Perl script removed bubbles that matched two sequences in the reference transcriptome and those with mapping ambiguity. The script also removed bubbles with a read coverage less than six and bubbles with one of the two alleles having less than two observations. Bubbles matching a transcript without a BLASTX hit were removed. These filtering steps ensured that sets of SNPs had the highest possible quality. The KisSplice/K2RT pipeline reports counts of both alleles at bi-allelic positions. Using these values, the distribution of allele frequencies at bi-allelic positions was plotted for each organism.

We tested the pipeline on RNA-seq data generated from different species with known ploidy or heterokaryon/homokaryon status to demonstrate the pipelines efficiency and robustness. The pipeline was applied on *R. irregularis* RNA-seq data to evaluate within-isolate polymorphism in RNA transcripts.

### Analysis of bubbles and bi-allelic transcripts

Transcript and bubble sequences were mapped to the reference genome with BLAT using the parameter -minIdentity=90. The alignments were parsed to quantify the proportion of uniquely mapped sequences.

To investigate whether bi-allelic positions discovered in RNA-seq data matched to bi-allelic sites in the genome, bubble sequences containing 2 alleles were mapped to the reference genome to provide the genomic coordinates of the bi-allelic positions. Next, allelic composition of each bi-allelic position in bubbles was compared to the allele composition of the position in the WG data. Any difference in allele composition was recorded as “inconsistent”. An “inconsistent” position is a bi-allelic position detected in RNA-seq that was not bi-allelic in WG data or that had a different allele composition to the WG data.

### Structural variation

WG samples (A1-WG4, A4-WG0, A4-WG9, A5-WG7, B3-WG6, C2-WG0, C2-WG3, DAOM-DNA1 and DAOM-DNA2), with a mean coverage greater than 20 were analysed to detect structural variations (SV). The calling of SV was performed using Lumpy version v2.16.2 [27]. This integrates multiple SV signals: read-depth variation events, paired-end events and split-read events. A new read mapping was performed with SpeedSeq version 0.1.2 [28]. All read sets were mapped to every genome to generate all possible combinations. The resulting .bam files were processed with Lumpy to call variants. Results were filtered to keep scaffolds with a size greater or equal to 10000, SV events that were supported by at least nine total events, three paired-end events and three split-read events. Only deletions detected with Lumpy were considered in this study.

SV calling was also performed using CNVnator [29]. The .bam files generated with novoalign to call the variants were reused to call the SV with CNVnator version 0.3.3. Results were filtered using the fthresholds: scaffold size greater or equal to 10000, p-value 1 (p-val1) < 0.05, q0 value < 0.5. Only deletions detected with CNVnator were considered in this study.

Genes overlapped by a deletion were found using BEDtools intersect. The script sma3_v2.pl was run on proteins deduced from genes to perform functional annotation and gene ontology enrichment analysis.

## Results

### Level of within-isolate polymorphism depends on the identity of the reference genome

The identity of the reference genome indeed had a strong impact on the density of poly-allelic positions (Figure S3), with the lowest density when ddRAD-seq data were mapped onto the respective genome assembly.

### Variants in ddRAD-seq and WG occurred at the same positions

The mean depth of coverage across the samples ranged from 22 to 33 and from 13 to 47 for ddRAD-seq and WG data, respectively (Figure S4). A greater number of predicted ddRAD-seq regions in the five isolates were covered by reads from WG data than by ddRAD-seq data (Figure S5). Poly-allelic positions were almost all bi-allelic. Tri-allelic positions represented approximately 0.5% of the poly-allelic sites. Only one tetra-allelic position was found (Table S5). Bi-allelic positions were found in all isolates and both types of data (Figure 2a). The density of bi-allelic positions ranged from 0.66 to 1.79 bi-allelic positions/kb (Figure 2b). In general, the density of bi-allelic positions was higher when the depth of coverage was higher (Figure 2b). The majority of bi-allelic sites that were found in the ddRAD-seq datasets also occurred in the WG datasets (Figure 3). In most cases, these occurred in all replicates of the two datasets or in several replicates of the ddRAD-seq data and WG data, confirming that it is very unlikely that they occurred because of random error (Figure 3). Hypergeometric tests showed that the overlap of which fragments from ddRAD-seq and WG mapped to the in silico predicted RAD fragments was highly unlikely to be due to chance (Cumulative hypergeometric test (function *phyper*), P < 0.00001, in all isolates). Alternative approaches to determining non-repeated regions yielded very similar results (Figures S6 and S7) meaning that the detection of within fungus SNPs was not strongly sensitive to the method of repeat detection. There were many more bi-allelic positions in the WG data that were not detected in the ddRAD-seq data simply because so many more predicted ddRAD-seq regions were covered by reads in WG data than in RD data (see Figure S5).

**Figure 2.**
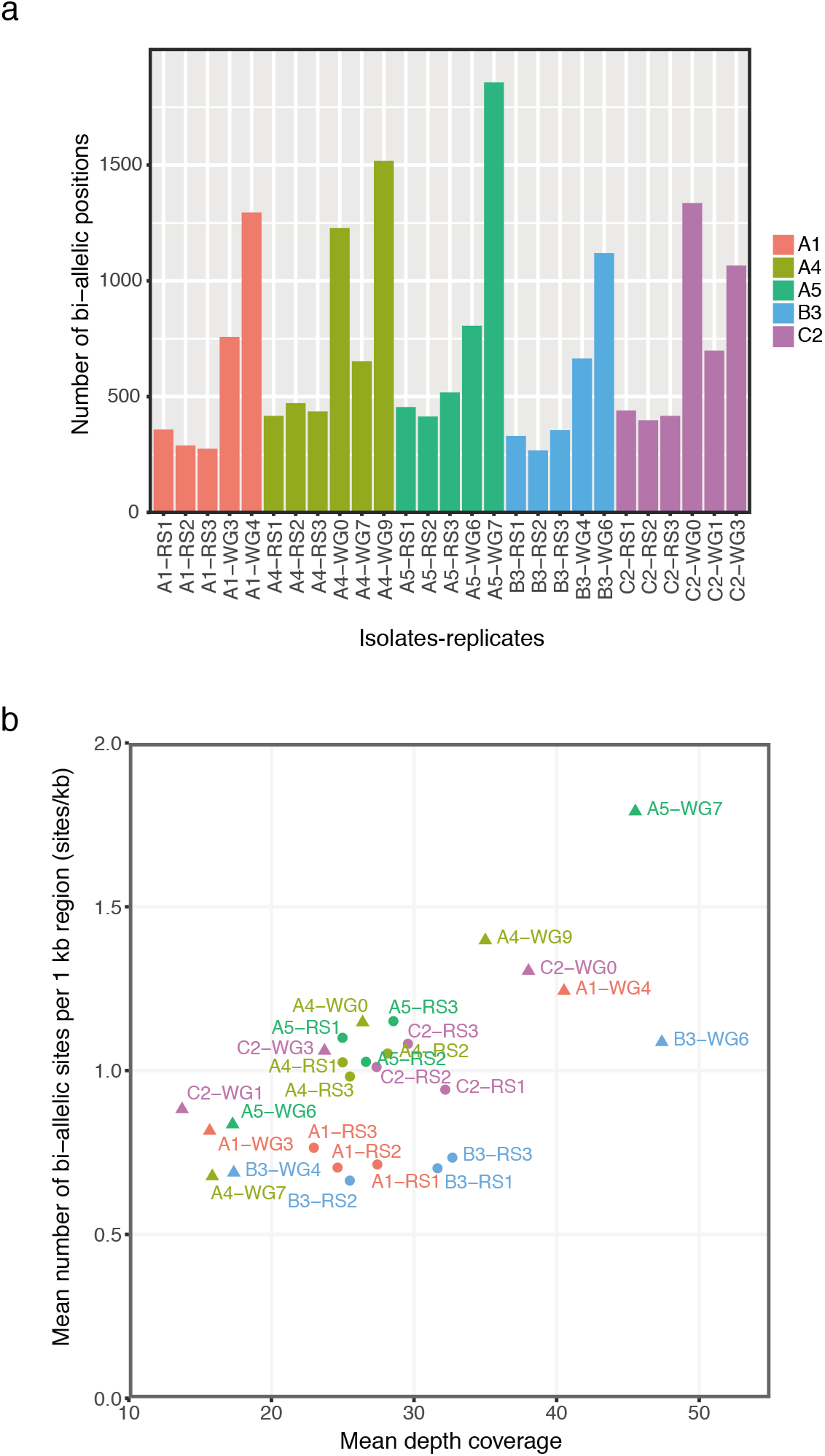
Detection of poly-allelic positions in whole genome (WG) and ddRAD-seq data. (a) Number of bi-allelic positions among replicate samples of the different isolates. (b) Mean density of poly-allelic positions (composed of SNPs) versus mean depth coverage (excluding ddRAD-seq loci with coverage lower than 10×) in predicted ddRAD-seq regions which are coding and nonrepeated (using method 03). “WGx” corresponds to the whole genome sequencing data, where “x” is the replicate identifier. “RSx” corresponds to the ddRAD-seq data, where “x” is the replicate identifier. Replicates of each isolate are shown as separate dots (RS) or triangle (WG).

**Figure 3.**
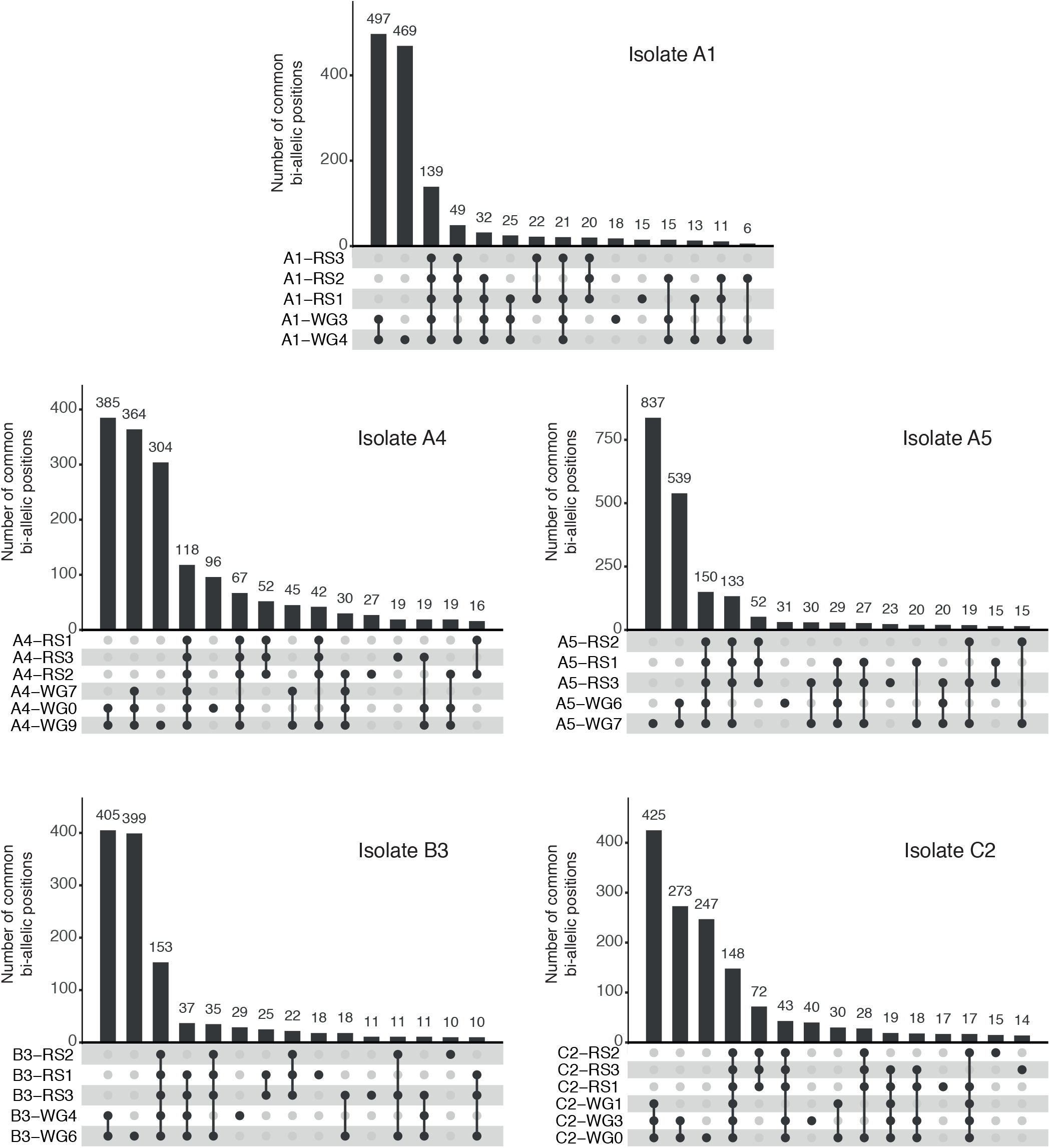
Comparison of bi-allelic sites found in replicates of WG and ddRAD-seq data in five *R. irregularis* isolates. Number of replicate samples from WG and ddRAD-seq data that contain the same bi-allelic sites in each isolate. The numbers above the bars in the histograms represent the number of bi-allelic sites shared among a given set of replicates from WG and ddRAD-seq. Bullets connected by vertical lines below the histograms show which replicates contained the same bi-allelic positions. The Up-set plots show numbers of common positions among samples ranked from highest on the left of the graph for the highest 15 combinations of samples. A very small number of bi-allelic sites were found among other samples that are not shown but these numbers are so low as not to alter the conclusions that can be drawn from this analysis.

### Allele frequency similarities

Assuming nuclei are haploid, allele frequency distribution at bi-allelic loci should reveal the number and proportion of different nuclei. Very similar allele frequency distributions were observed among the replicates of each isolate originating from WG and ddRAD-seq data (Figure 4; Figure S8). A peak at 50:50 was detected in A4 and A5 (Figure 2b), as expected for a 50:50 heterokaryon and as shown previously [9]. There was a slight peak at 50:50, as well as slight peaks at approximately 20:80 and 80:20, in A1, B3 and C2 (Figure 4). It is also noticeable that the small peaks at 20:80 and 80:20 also existed in the isolates A4 and A5 despite a clear peak at 50:50. The isolate DAOM197198 was also subjected to the same analysis. These results are presented in the Figure S9.

**Figure 4.**
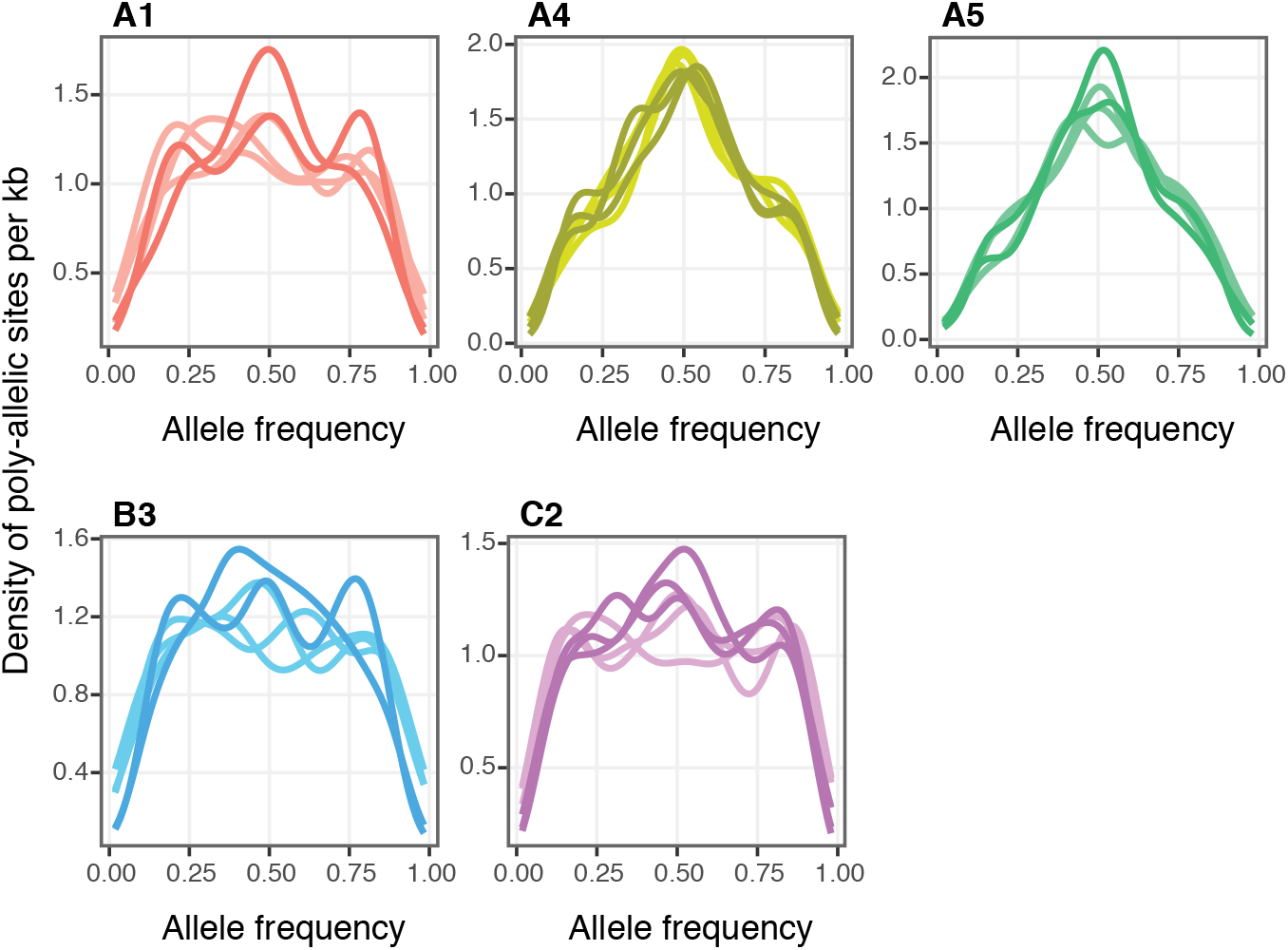
Comparison of distribution of allele frequencies in five *R. irregularis* isolates observed in WG and ddRAD-seq data. Distribution of allele frequencies at poly-allelic positions (composed of SNPs) in non-repeated and coding regions (using method 03) of the isolates A1, A4, A5, B3 and C2. Dark coloured lines represent WG data and clear lines ddRAD-seq data.

### Effects of coverage and other parameters

We investigated how different parameters can affect the discovery of bi-allelic positions. All highest-covered samples had a depth of coverage lower than 50× (Figure S4). In Figure 2b, the density of bi-allelic positions was only reported in non-repeated, coding regions of predicted ddRAD-seq fragments and was higher when the depth of coverage was higher. The number of bi-allelic positions was measured in the whole genome and not only in the predicted ddRAD-seq regions (Figure S10). The greatest numbers of poly-allelic positions were found in samples having the highest coverage depth. Increasing the depth of coverage by sequencing more deeply lead to higher number of bi-allelic positions and no saturation was detected (Figure S10).

We applied different thresholds on the depth of coverage in order to evaluate the effects of coverage on the discovery of bi-allelic positions and on patterns of allele frequencies (Figure S11). Increasing the coverage thresholds had a strong impact on allele frequency distribution. At higher thresholds, the signal around 50:50 disappeared, including the strong 50:50 peaks found for A5 and A4. High thresholds led to a U-shape distribution similar to what was observed by Ropars et al. (2016) [9].

### Bi-allelic positions within RNA transcripts

We used RNA-seq data to explore whether within-isolate polymorphism was detectable in transcripts. We first tested the pipeline on data generated from different species for which the ploidy level is known or for which the states homokaryon, heterokaryon, homozygote and heterozygote is already known. The pipeline clearly demonstrated its efficiency to distinguish heterozygotes from homozygotes, and heretokaryons from homokaryons (Figure S12 and Supplementary information).

The number of transcripts assembled independently from the reference genome was greater in the isolate C3 than in the isolates C2 and DAOM197198 when no filtration was applied (Figure 5a). However, the number of genes deduced from Trinity-assembled transcripts was very similar in the three isolates (Figure 5a). The number of genes recovered in the transcripts represented 71%, 77%, and 71% of the 22 812 genes that are thought to be present in the *R. irregularis* genome in C2, C3 and DAOM 197198, respectively. This meant that the RNA-seq datasets from these isolates should have allowed the recovery of a significant proportion of the bi-allelic transcripts, if they exist. Transcripts with bi-allelic positions were found in all isolates (Figure 5b), with a greater proportion in C3 than in C2 and DAOM197198. Detailed results can be found at https://github.com/FredMasc/PAAT2019. The percentage of bi-allelic sites with synonymous codons was 54%, 41% and 37% in C3, C2 and DAOM, respectively.

**Figure 5.**
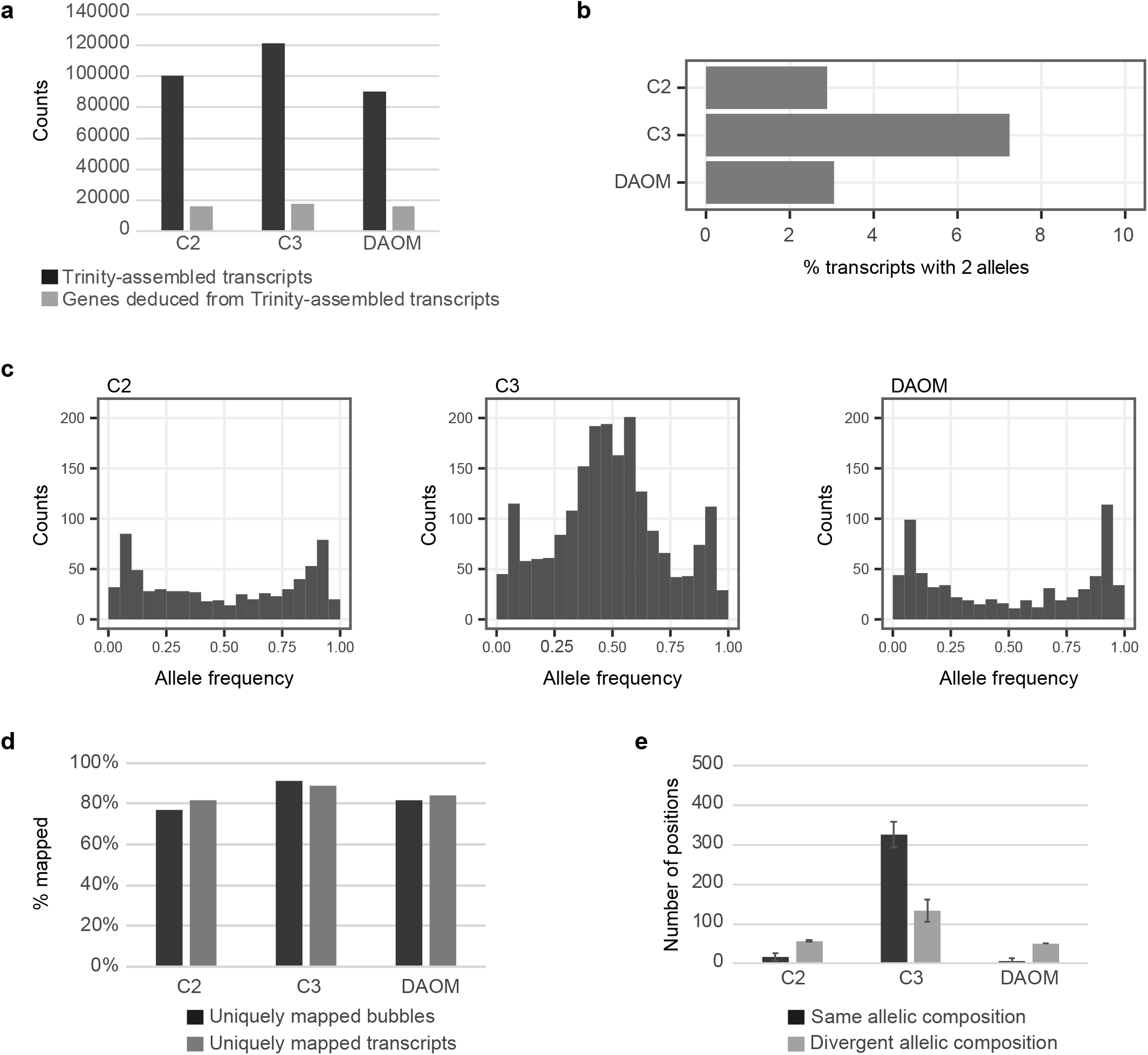
Detection of bi-allelic positions in RNA-seq data. (a) Number of transcripts (black) and genes (grey) obtained with the de-novo transcript assembly approach. (b) Percentage of transcripts containing bi-allelic positions in isolates C2, C3 and DAOM197198. (c) Allele frequency graphs of RNA-seq data at bi-allelic sites. (d) Mapping percentages of transcripts (grey) and bubbles (black) on reference assemblies. Sets of bubbles and transcripts are mapped on the appropriate assembly for each isolate. (e) Number of bi-allelic positions detected in RNA-seq data that have the same allele composition (black) or divergent allele composition (light grey) compared to WG data. Error bars represent ± 1S.E.

Allele frequencies in C3 displayed a peak centred at 50:50, although the peak was not as clear as in some of the diploid or heterokaryons shown in Figure S12. There were two additional peaks at 10 and 90. C2 and DAOM197198 allele frequency plots did not exhibit a 50:50 peak (Figure 5c). U-shaped distributions were observed in C2 and DAOM197198. In addition, moderate peaks were observed at 10:90 and 90:10 in the three isolates. Such patterns are very similar to the patterns observed in the ddRAD-seq and WG data (Figure 4 and S10).

### Testing for the existence of matching bi-alleles in the *R. irregularis* transcriptome and genome

We tested whether bi-alleles found in the transcriptome of the three isolates were also detected independently in the genome. Moderate peaks at 0.1 and 0.9 in the frequency plots (Figure 5c) could be due to wrongly called SNPs because pairs of bubbles do not belong to single transcripts. We tested if all the bubble sequences produced by KisSplice mapped unambiguously to their corresponding reference genome. Both bubble and whole transcript sequences showed similar, and high, proportions of transcripts mapping a single time to the reference genome (Figure 5d). Second, these peaks could be due to sequencing errors. We tested if the positions with two alleles found in RNA-seq data also displayed the same two alleles in WG data. If the bi-alleles are located on two genetically-different nuclei then the same alleles should be found in RNA-seq data and WG data. A total of 326 identical bi-allelic positions occurred in both RNA-seq and WG data in the isolate C3 (Figure 5e). A very small number of bi-allelic positions found in RNA-seq data were also found in WG data in C2 and DAOM197198 (17 in C2 and 5 in DAOM197198). Most of the bi-allelic positions detected in RNA-seq data in C2 and DAOM197198 were not found in WG data (depicted by grey bars and labelled as “divergent allele composition”; Figure 5e). An example of inconsistent positions in C2 is provided in Figure S13. As a comparison, an example of consistent positions in C3 is shown in Figure S14. A notably larger number of bi-allelic positions occurred in RNA-seq data that were not found in WG data in isolate C3 compared to C2 and DAOM197198 (Figure 5e).

### Impact of unsolved regions on the within-isolate polymorphism

Each position was visually inspected in the WG read alignment to the reference genome. Bi-allelic positions in C2 and DAOM197198 were frequently found around problematic regions in the genome assembly. Problematic regions were either unsolved regions (gaps) in the genome assembly or regions where the depth of coverage shows an unexpected increase compared to the mean depth of coverage. Both types of problematic regions are frequently found in close locations. Unsolved regions are regions in which Ns are reported in the middle or at the extremities of the scaffold sequences. They correspond to genomic regions that are difficult to assemble. The unsolved regions, which are likely to occur in repeated regions, led to the reads to map wrongly and to accumulate at the wrong locations. This transfer of reads generated bi-allelic positions containing frequently two alleles with equal proportions. Examples of unsolved regions and their effect on the detected within-isolate polymorphism are provided in Figure S15.

### Structural variation in the genome of the three isolates

Within isolate variation could also be potentially caused by structural variation existing among nuclei. Structural variation was found in all isolates. Both CNVnator and Lumpy enabled us to detect deletions in all isolates (Figure 6a and 6b). More deletions were detected in C3 than in C2 and DAOM197198 with both methods. The number of deletions detected was greater with CNVnator than with Lumpy, but the pattern was very similar. Most deletions were smaller than 5 kbp (Figure 6c). The largest sized deletions (ranging between 15 and 20 Kbp) were found in the genome of A4, but their number was low and could not be linked to the enrichment of a particular group of genes of similar function. Genes that occurred where a deletion had also occurred were identified and their number is shown in Figure 6d. Detailed results can be found at https://github.com/FredMasc/PAAT2019. Functional annotation and GO enrichment analysis of these genes did not reveal large differences among the isolates (Table S6). Notably, the GO category “DNA binding” was prevalent and many genes matched with transposases. Visual inspection of the alignments was conducted to characterize the deletions. Many deletions appeared to be associated with unsolved regions. The assembly quality is an important parameter to accurately detect deletions using Illumina small read mapping. We looked for genes with functions unrelated to transposition. Regions where structural variants were found in all isolates with a strong drop in coverage were discarded because they likely reflect problems in genome assembly. We provide two examples of genes in A4/C3 that overlap deletions that did not occur in the other isolates (Figure 6e and 6f). The region around each gene was affected by a decrease of approximately 50% in coverage. In addition, SV at gene *g11881* (Figure 6e) was supported by 41 read pairs that had an abnormal insert size (too large) and SV at gene *g22991* (Figure 4f) was supported by 19 read pairs that had excessively large insert size. The gene *g11881* encodes a hypothetical protein (148 amino acids) with no obviously identifiable domain. The gene *g22991* encodes a serine/threonine protein kinase (487 amino acids).

**Figure 6.**
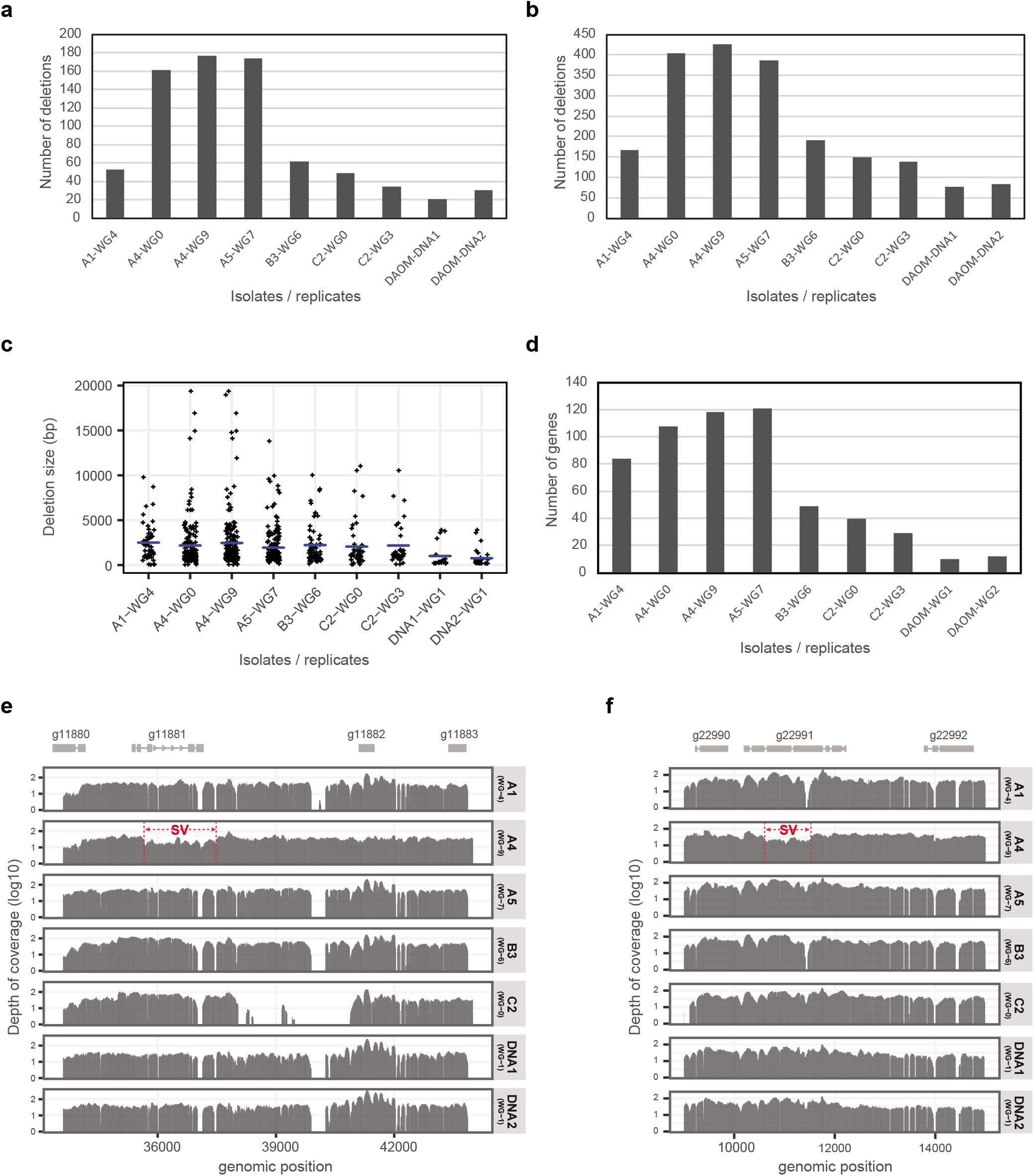
Structural variation (SV) in WG data. (a) Number of deletions found with Lumpy. (b) Number of deletions found with CNVnator. (c) Deletion sizes among the isolates. (d) Number of genes overlapped by deletions. (e) and (f) Examples of structural variation found in A4 at the gene *g11881* (e) and at the gene *g22991* (f). The first track represents the gene model. The other tracks correspond to the depth of coverage measured in different isolates. The genomic region affected by an SV is labelled with a red box and red arrows.

## Discussion

Here we demonstrated that studies of within and among-isolate SNP variation in *R. irregularis*, conducted independently, and using different techniques produced remarkably similar results. We have also made the first analysis linking within-AMF isolate variation at the genomic level to within-fungus transcription at bi-allelic sites indicating that in a heterokaryon *R. irregularis*, transcription of two alleles at bi-allelic sites is consistent with the proportion of the two genetically different nuclei but that in homokaryons little evidence exists for any expression of possible within fungus genetic variation. Finally, we have demonstrated the level of structural variation in the genome in 3 isolates that was due to deletions. The implications of these findings are discussed below.

### Within and among-isolate SNP variation in *R. irregularis*

Our analyses indicated that SNP variation within isolates of *R. irregularis* were remarkably similar in two studies where the same fungal isolates (or clones of the isolates) were cultured and then sequenced independently in two different laboratories and using two different sequencing approaches. Not only were many of the same SNPs identified in the two studies but they were repeatedly observed in replicates within and between the studies (Figure 3). The same bi-allelic SNPs observed both among replicates and between independent studies means that the probability that they occurred in the sequence datasets by chance is exceptionally unlikely. Furthermore, allele frequency plots among samples from the two studies are also remarkably similar (Figure 4). This analysis shows that for detection of SNPs in *R. irregularis*, both approaches are equally valuable for SNP discovery in this fungal species, with the only differences being that more SNPs can be discovered with whole genome sequencing but more isolates can be screened for the same cost using ddRAD-seq. In both cases, sufficient sequencing depth is required and the fact that deeper sequencing reveals more SNPs means that deep sequencing might lead to discovery of a greater number of rare alleles. However, this should also be coupled with replication to ensure rare alleles truly exist.

The discrepancies between the studies are that Ropars et al. (2016) reported lower within-fungus polymorphism than Wyss et al. (2016), although this differed greatly among isolates with values for some isolates being very close and others more distant [8, 9]. As shown in this study, one reason for this is that Wyss et al. (2016) mapped reads to the only available reference genome at that time; namely, an assembly of the isolate DAOM 197198. This is because of the very large amount of among-isolate genetic variation. Mapping to a genome assembly of the isolate rather than a reference genome greatly reduced the number of bi-allelic SNPs identified (Figure S3). This is because of the very large genetic differences among isolates in this species. This is an important result as looking at SNPs among individuals in population genetics studies is usually conducted by comparing SNPs found in the genome or transcriptome of a given individual to a reference genome. However, artefacts arising from mapping to one reference genome still does not account for the discovery of consistent among replicate or between-study within-fungus variation observed in each isolate. In the case of isolates considered heterokaryons, these likely represent differences between the two nucleus genotypes or some artefact of mapping where problematic parts of the genome assembly occur. In isolates considered homokaryons, bi-allelic SNPs could either represent artefacts due to problematic parts of the genome assembly or rare alleles.

Allele frequency plots of some isolates reported in Wyss et al. (2016) did not exhibit the U-shaped curves [8]. However, Ropars et al. (2016) observed U-shaped curves in homokaryons but did not explain what accounted for such a distribution [9]. In this study, we show that the shape of the allele frequency distribution is highly sensitive to the read threshold number used as a cut-off (Figure S11). At a low threshold of 10, almost all isolates appeared to have multiple bi-alleles with many different proportions; even those claimed to be homokaryons. It is difficult to set a standard threshold for all isolates at which homokarons appear as homokaryons and heterokaryons reveal a 0.5 peak of allele distribution. A higher threshold sets a bias for where only high coverage variants are retained and these usually represent areas of poor assembly or repeated regions. Thus, this represents and artefact where even heterokaryons will exhibit a U-shaped curve of allele distribution making it difficult to detect heterokaryons. Given that setting a read threshold number is arbitrary, this makes distinction between homokaryons and heterokaryons difficult.

### Bi-allelic variation in the *R. irregularis* genome and its transcription

While there has been considerable effort in documenting within and among isolate genome variation in AMF, little attention has been paid whether this variation is transcribed. The number of bi-allelic transcripts that were observed in the isolates expected to be homokaryons were indeed low and only a very small number matched those observed in the genome SNP data (Figure 5e). Of those, most mapped to problematic regions of the genome assembly indicating that these are probably artefacts. A notably large number of alleles in the heterokaryon C3 were observed as 2 alleles in the transcript data and in the genome. We assumed that bi-allelic transcripts that did not have corresponding alleles in the genome data were artefacts (light grey bars in Figure 5e). However, these were notably higher in C3 than in the other two isolates which would not be expected if they occurred because of sequencing errors. Some bi-allelic positions were not discovered in the genome of this isolate because the coverage of genome data is insufficient. The results show that indeed, genes of the two different nucleus genotypes present in the heterokaryon C3 are transcribed. About 50% of these bi-alleles in C3 contained non-synonymous substitutions meaning that this variation among nuclei could potentially affect the fungal phenotype. The allele frequency graphs of bi-allelic SNPs in the genome and those observed independently in the transcriptome both revealed an allele frequency of 0.5. This suggests that in isolate C3, the two nucleus genotypes are present in roughly equal proportions; a measurement that was also corroborated by a separate study [30] using the allele frequencies of a single marker and replicate measurements on the same isolate. The 0.5 allele frequency in the transcripts suggests that allele expression from a pair of given alleles located on the two nuclei are expressed in equal amounts from two differing parts of the two nucleus genomes. In several studies from our group, sibling cultures of AMF, initiated from single sibling spores for the parent C3 have been shown to exhibit quantitative traits such as hyphal growth and spore production, as well as their effects on plant growth, that were significantly different from the parent C3 [4, 5]. This is highly unusual for clonal offspring. Recently, the ratios of the two nucleus genotypes in some of these sibling offspring lines were shown to deviate significantly from the 50:50 ratio observed in the parent C3 [30]. If gene transcription from the two nucleus genotypes is proportional to the frequency of those genotypes (as shown in the present study) then we could expect that sibling clonal offspring of C3 could have unequal transcription of bi-alleles and this could account for the differences in quantitative traits observed among single spore clonally produced siblings of C3.

### Structural variation in homokaryon and heterokaryon *R. irregularis*

The analysis of structural variation in the isolates revealed a higher level of structural variation in the dikaryon C3 than the other two isolates. If C2 and DAOM197198 are truly homokaryons then this structural variation would not be expected. Thus, we assume that this variation is also due to parts of the assembly where data is of poor quality. In C3, the higher level of structural variation should represent true structural differences occurring between the two different nucleus genotypes and such structural variation among the two nucleus genotypes could also have a regulatory effect influencing the fungal phenotype. These structural variations could either have arisen because of the fusion of two homokarons carrying genetically different nuclei or by rearrangement and recombination among nuclei within C3, as reported previously [31].

### Conclusions

Investigating whether within-AMF isolate genetic variation is extremely difficult. Independent studies using different techniques have identified the same variants and we show that this has obviously not occurred by chance. Transcription profiling is a good way to verify some of these bi-allelic positions in the genome. Transcription profiling did not reveal significant transcription of bi-allelic sites in a homokaryon isolate. However, transcription profiling reveals that indeed, within fungus genetic variation existing in heterokaryons is transcribed from both of the co-existing genomes and in equal proportions from both nucleus genotypes. Further studies are needed to determine the nature of the genetic differences between these two genotypes and how their expression influences the fungal phenotype and its effects on plants.

## Acknowledgements

We thank Vincent Lacroix for his advice on KisSplice and the Genomic Technologies Facility, University of Lausanne for sequencing. Bioinformatics computations were performed at the Vital-IT (www.vital-it.ch) Centre for High Performance Computing, Swiss Institute of Bioinformatics. This research was funded Swiss National Science Foundation grant (number: 31003A_162549).

## Author Contribution

Design of the research: FGM, TW, MP, IRS; material and sample preparation: TW, PR; data analysis and interpretation: FGM, TW, MP, IRS; writing the manuscript: FGM, IRS, TW.

