## Supplemental methods, results & figures for "Investigating unexplained genetic variation and its expression in the arbuscular mycorrhizal fungus *Rhizophagus irregularis*" (from another version of this preprint: version 1, 2019-06-26)

### Supplementary results

**Test of the pipeline including KisSplice on RNA-seq data generated from different species**

We investigated the level of within-isolate polymorphism in transcripts of two homokaryote and one dikaryote isolate of the fungus *Antrodia cinnamomea*. More transcripts with poly-allelic positions were found in the dikaryote than in the homokaryotes (Figure S12a). The allele frequency graphs showed a peak at an allele ratio of 50:50 in the dikaryon (Figure S12a). It is also the typical peak expected for a dikaryon having an equal ratio of different nuclei that equally express transcripts. Only a weak random signal, considered as artefactual, was observed in the allele frequency graphs of the homokaryons (Figure S12a). The dikaryotes could be unambiguously identified among the homokaryotes and the allelic transcripts are expressed, on average, at the same level.

The pipeline was also tested with four *Arabidopsis* RNA-seq samples. Three of the samples were natural diploid accessions and mostly homozygous as *Arabidopsis* is self-fertilizing. The fourth sample was a lab-generated F1 hybrid between two different natural accessions and can be considered as a strong heterozygote. Analysis of the allele frequency graphs clearly showed an artefactual signal for the three natural accessions and a high 50:50 peak for the hybrid (Figure S12b). The heterozygote could be unambiguously identified among the homozygotes and the allelic transcripts are expressed on average at the same level.

RNA-seq data from other plants with different levels of ploidy were also subjected to the pipeline. *Aegilops longissimi* is a diploid homozygote that was hybridized with another diploid homozygote (*Triticum urartu*) to create the heterozygote tetraploid hybrid. Only an artefactual signal was found for the homozygote *A. longissimi*, while a high 50:50 peak was found for the heterozygote tetraploid hybrid plant (Figure S12c). The heterozygote could be unambiguously distinguished from the homozygote and the transcripts of bi-alleles were expressed, on average, at the same level.

Analysis of RNA-seq data from the haploid *Saccharomyces cerevisiae*, revealed no obvious peak, as expected. The heterokaryote *Rhizoctonia solani* exhibited a 50:50 allele frequency peak, although some isolates of this species have been reported to be heterokaryotic and harbouring more than 2 genetically different nuclei. If this was the case in this study then the pipeline was not able to resolve more complex within-fungus genetic structure. The diploid *Candida albicans* exhibited an expected 50:50 peak (Figure S12d). Allelic transcripts in the heterokaryote and in the diploid were expressed, on average, in similar proportions.

In all the tested RNA-seq data from different species the pipeline, including KisSplice, demonstrated its efficiency to distinguish the heterozygotes (and heterokaryotes) from the homozygotes (and homokaryotes) without ambiguity.

### Supplementary material and methods

##### Versions, flags and options used for bioinformatics programs

**Genomics**

tagcleaner.pl version 0.14

```
    -tag5 CCTACACGACGCTCTTCCGATCT          (forward reads)
    -tag3 AGATCGGAAGAGCACACGTCTGA
    -matrix exact
    -trim_within 60
    -mm5 3
    -mm3 4

    -tag5 TCAGACGTGTGCTCTTCCGATCT          (reverse reads)
    -tag3 AGATCGGAAGAGCGTCGTGTAGG
```

prinSeq-lite.pl version 0.20.4

-trim_ns_left 1 -trim_qual_type "mean" -trim_qual_window 10 -trim_qual_step 9 -trim_qual_right 15 -min_len 50

process_radtags version 1.06 (STACKS)
-E phred33 -r -e ecoRI -c -q -D -s 10

Novoalign 3.02.12
Standard parameters

samtools version 1.3
Standard parameters

freebayes version 1.0.2
-p 10 -J -K -F 0.1 -0 -u

vcffilter 1.0
-f 'QUAL > 30'

RepeatModeler Open-1.0
Standard parameters

RepeatMasker Open-3.0
Standard parameters

fasta-36.3.5e
Standard parameters

**Transcriptomics**
Assemblies with Trinity
Trinity version 2.5.1
bowtie2 version 2.3.0
java version 1.8.0_121
Trinity
--trimmomatic --max_memory 100G --CPU 60 --SS_lib_type RF --full_cleanup

kissplice version 2.4.0
--experimental --max-memory 6000 -v -s 2 -k 41 -t 10

ncbi-blast version 2.6.0+
Standard parameters

TransDecoder-v5.0.2
Standard parameters

Blat v36
-minIdentity=80

**Structural variations**

Speedseq v0.1.2

Standard parameters (as specified in LUMPY manual)

lumpyexpress (LUMPY 0.2.13)

Standard parameters

SVTyper 0.7.0

Standard parameters

CNVnator 0.3.1

Standard parameters

Subsequent filter for scaffold size (10000), p-value n°1 (< 0.05), q0 value (< 0.5)

#### Custom Perl and R scripts

Custom Perl and R scripts used to treat the data in Genomics, Transcriptomics and Structural variations are availalble at https://github.com/FredMasc/PAAT2019

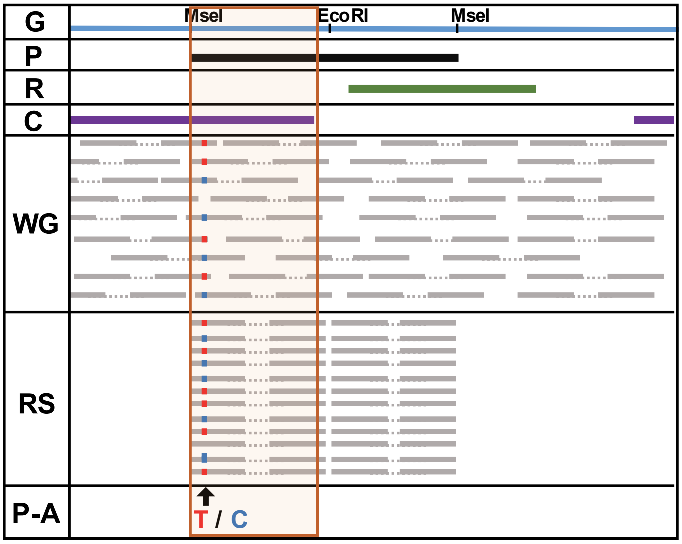


**Figure S1. Methodology to detect poly-allelic positions in whole genome (WG) and ddRAD-seq (RS) data.** Schematic diagram of the methodology used to discover potential within-fungus genetic polymorphism. Line 'G' corresponds to a linear part of a genome where EcoRI and MseI restriction sites are marked. Line 'P' shows a black bar corresponding to a predicted RAD fragment between EcoRI and MseI restriction sites. Line 'R' shows a green bar corresponding to a repeated region in the genome. Line 'C' shows a violet bar corresponding to a coding region in the genome. Lines 'WG' and 'RS' show respectively whole genome sequencing reads and ddRAD-seq reads mapped to this genomic region. Line 'P-A' shows the position where poly-allelic positions are found and the alleles found at this position. A semi-transparent orange box delimits the region of interest: in the predicted RAD fragment, in coding region and not in the repeated region. The example shows a position where alleles T and C can be found. In the empirical data, most poly-allelic positions are bi-allelic positions.

a

b

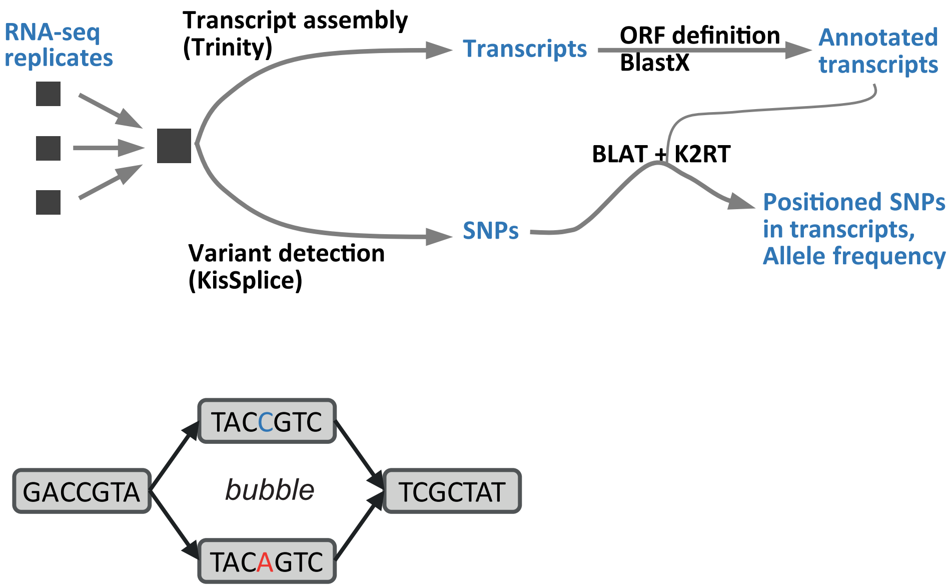


**Figure S2. Analysis pipeline of RNA-seq data.** (a) Overview of the pipeline including KisSplice to call SNPs from RNA-seq data without a reference genome. (b) Example of a 'bubble' structure shaped in a De Bruijn graph at a given bi-allelic position.

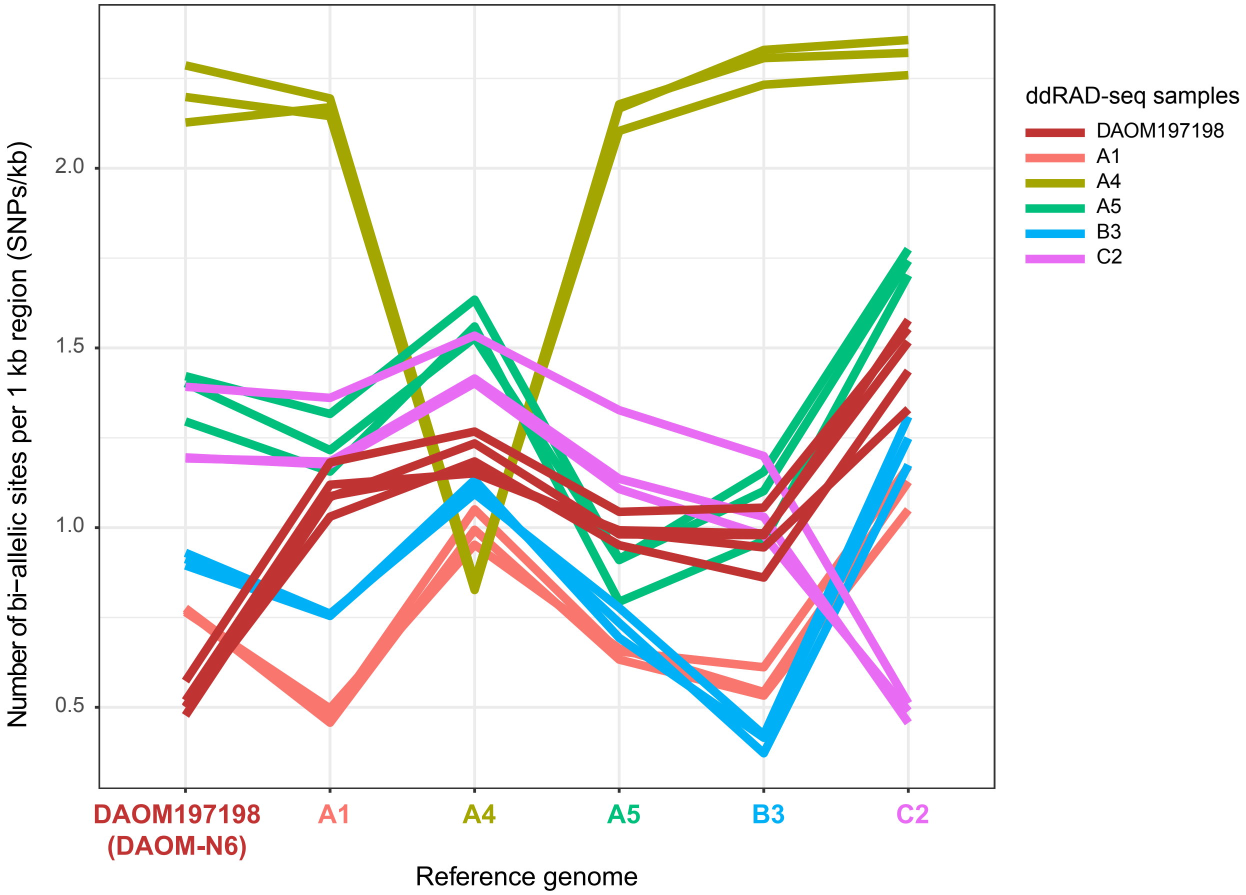


**Figure S3: Effects of the reference genome on the density of bi-allelic positions.**
Each line represents the density of bi-allelic positions detected in one replicate when ddRAD-seq reads were mapped to different reference genomes. A specific colour is used for each isolate.

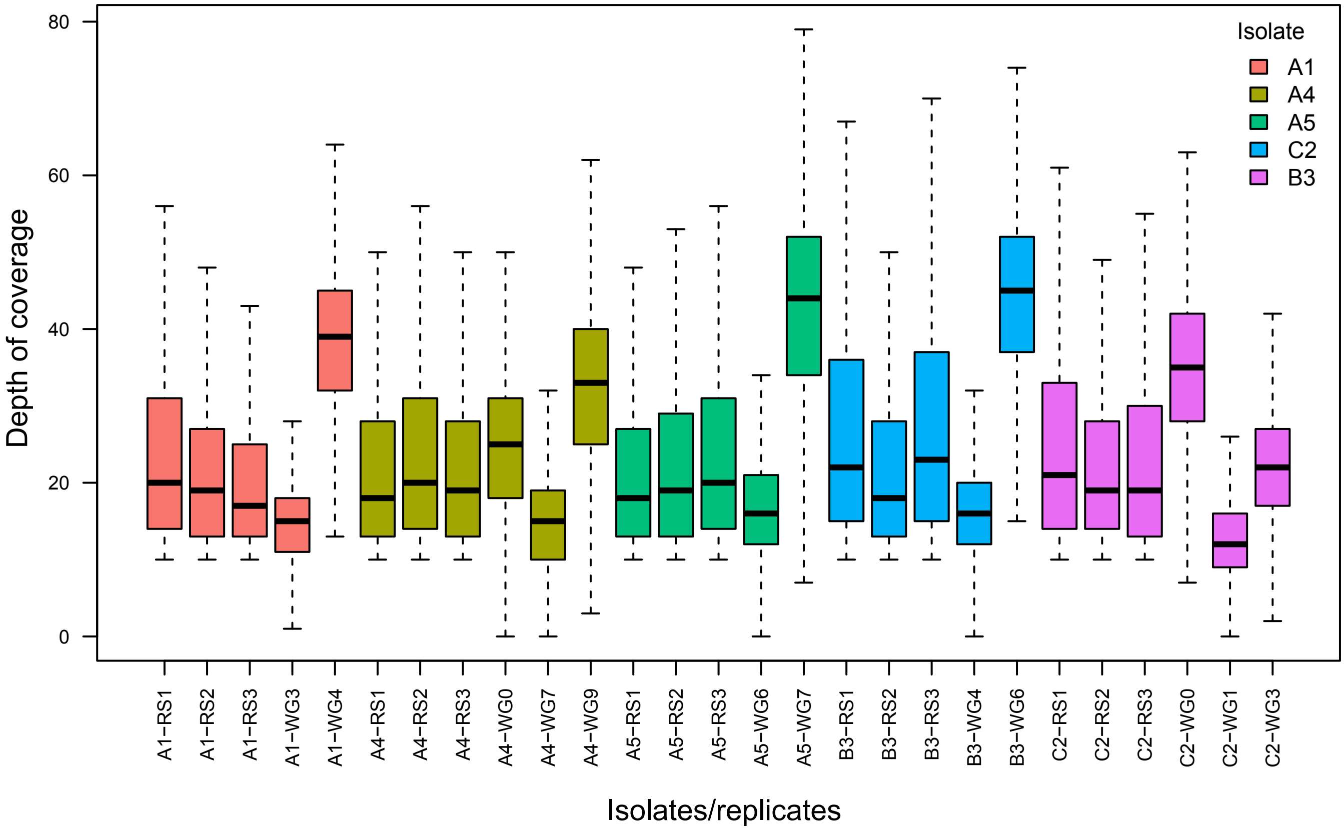


**Figure S4: Boxplot of the depth of coverage of ddRad-seq (RS) data and whole genome sequencing (WG) data.** All samples were aligned to their respective genomes. The boxplot depicts the distribution of depth of coverage of all ddRAD-seq and whole-genome sequencing samples. Artifactual sequences present in the ddRAD-seq data were eliminated by applying a minimal coverage threshold of 10×.

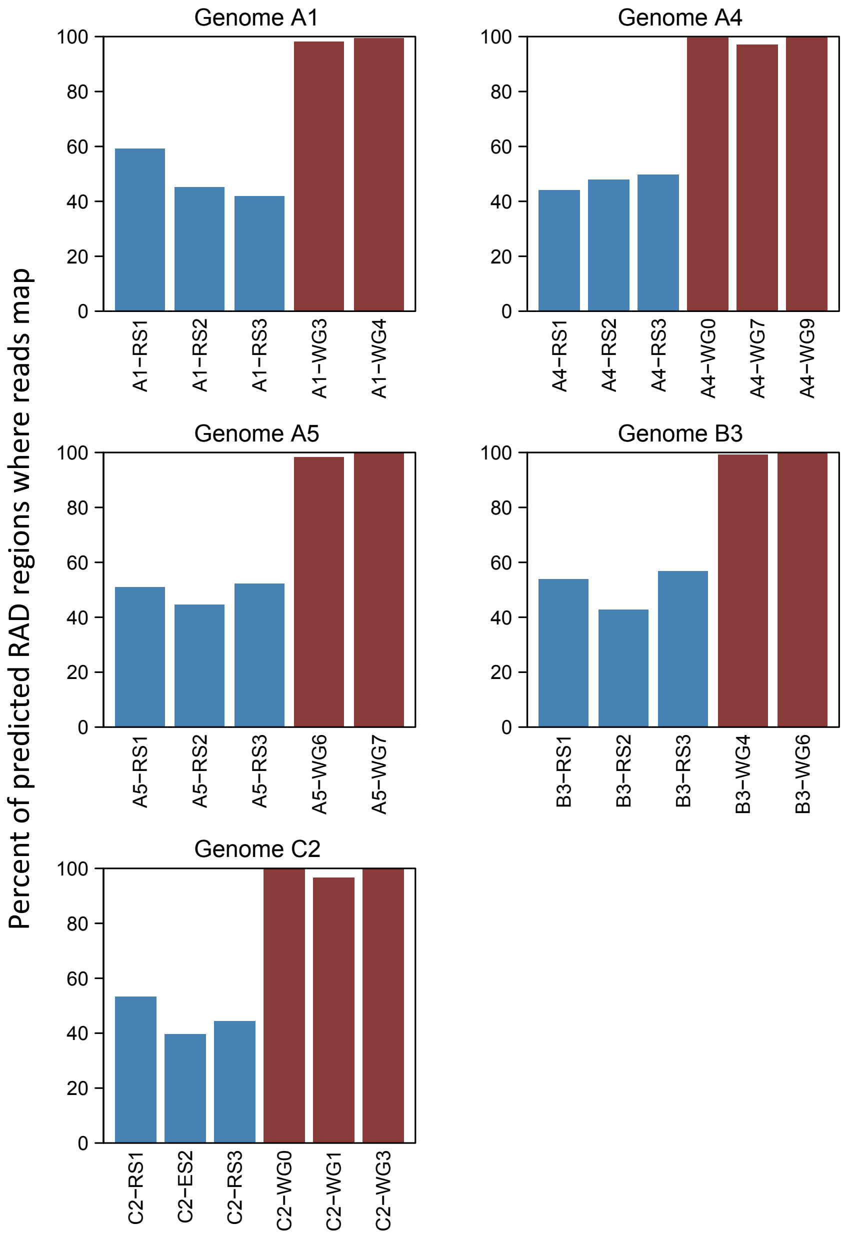


**Figure S5: Percentage of predicted ddRAD-seq regions covered with ddRAD-seq reads and whole genome sequencing (WG) reads.** Blue bars correspond to ddRAD-seq data and red bars correspond to WG data. Only ddRAD-seq regions with a depth coverage threshold of 10× were recorded. Graphs were generated in non-repeated and coding ddRAD-seq regions, following the M03 method to identify the repeats.

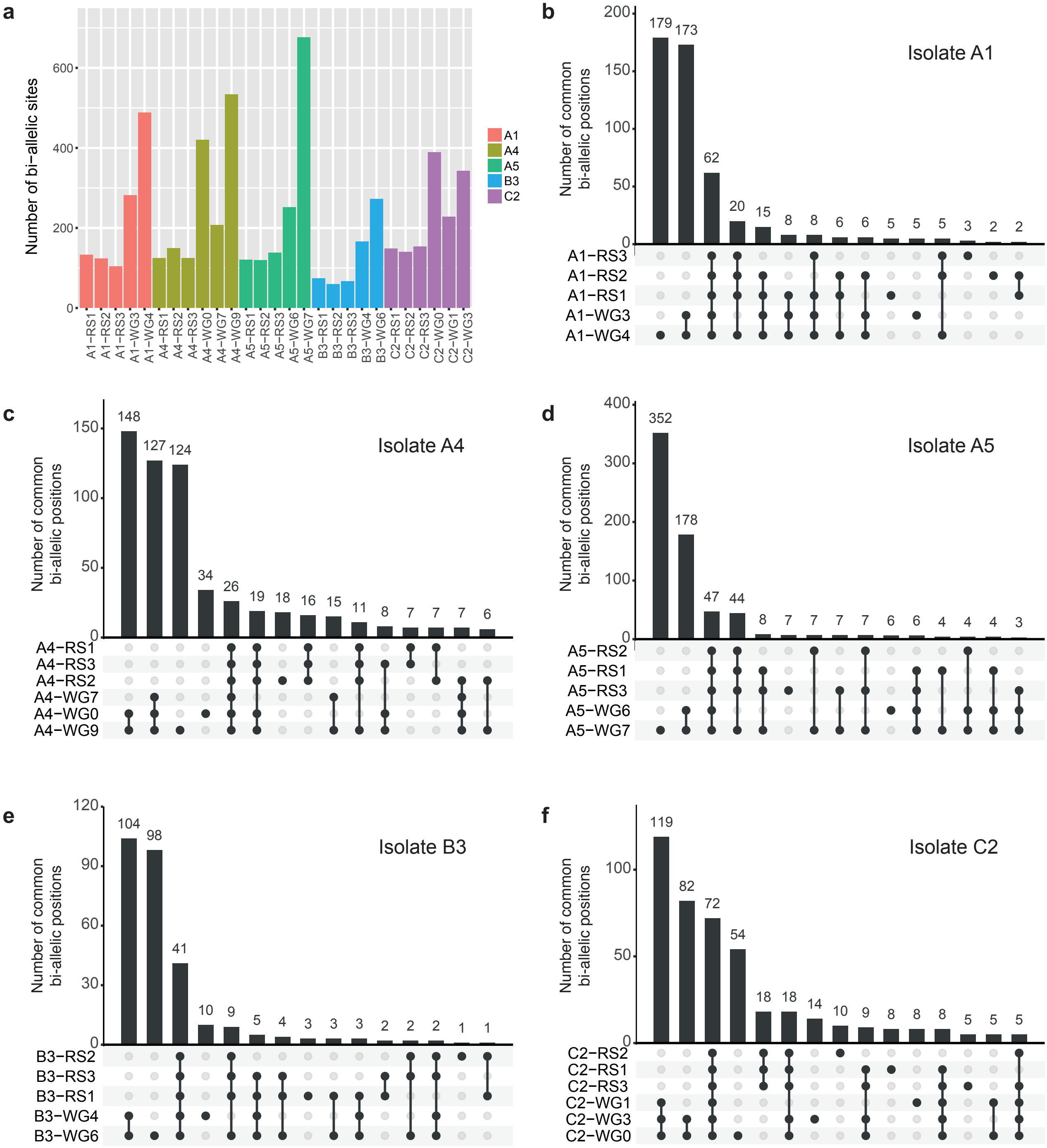


**Figure S6: Number of common bi-allelic positions in non-repeated (defined with method 07) and coding genomic regions among samples in different isolates of *R. irregularis*.** (a) Number of bi−allelic positions among replicate samples of different isolates. (b), (c), (d), (e) and (f) Number of common bi−allelic positions among samples of A1, A4, A5, B3 and C2 respectively. Numbers of common bi−allelic positions are shown with UpSet plots. The vertical lines connecting bullets show common bi−allelic positions in each set of samples.

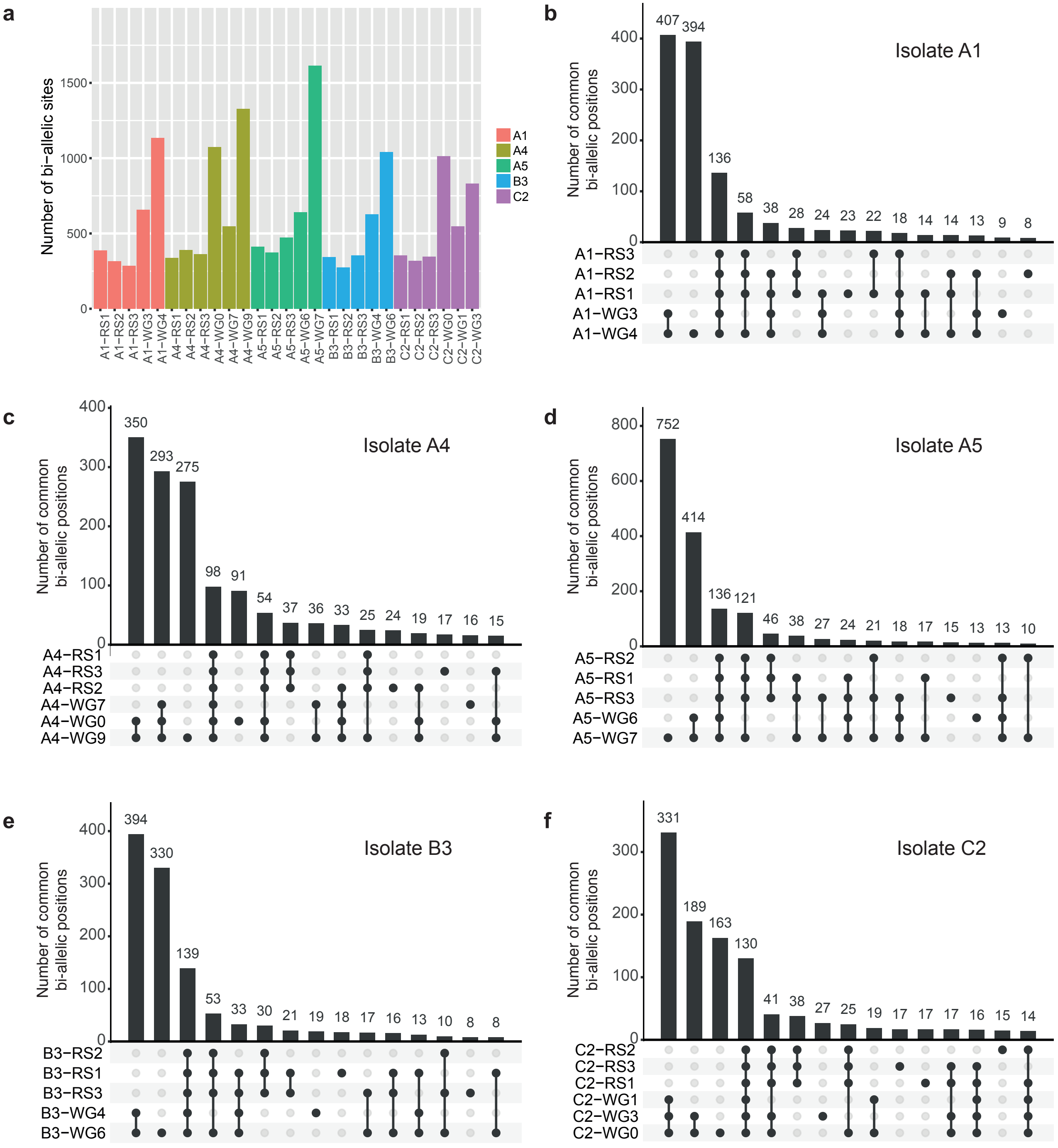


**Figure S7: Number of common bi-allelic positions in non-repeated (defined with method 12) and coding genomic regions among samples in different isolates of *R. irregularis*.** (a) Number of bi−allelic positions among replicate samples of different isolates. (b), (c), (d), (e) and (f) Number of common bi−allelic positions among samples of A1, A4, A5, B3 and C2 respectively. Numbers of common bi−allelic positions are shown with UpSet plots. The vertical lines connecting bullets show common bi−allelic positions in each set of samples.

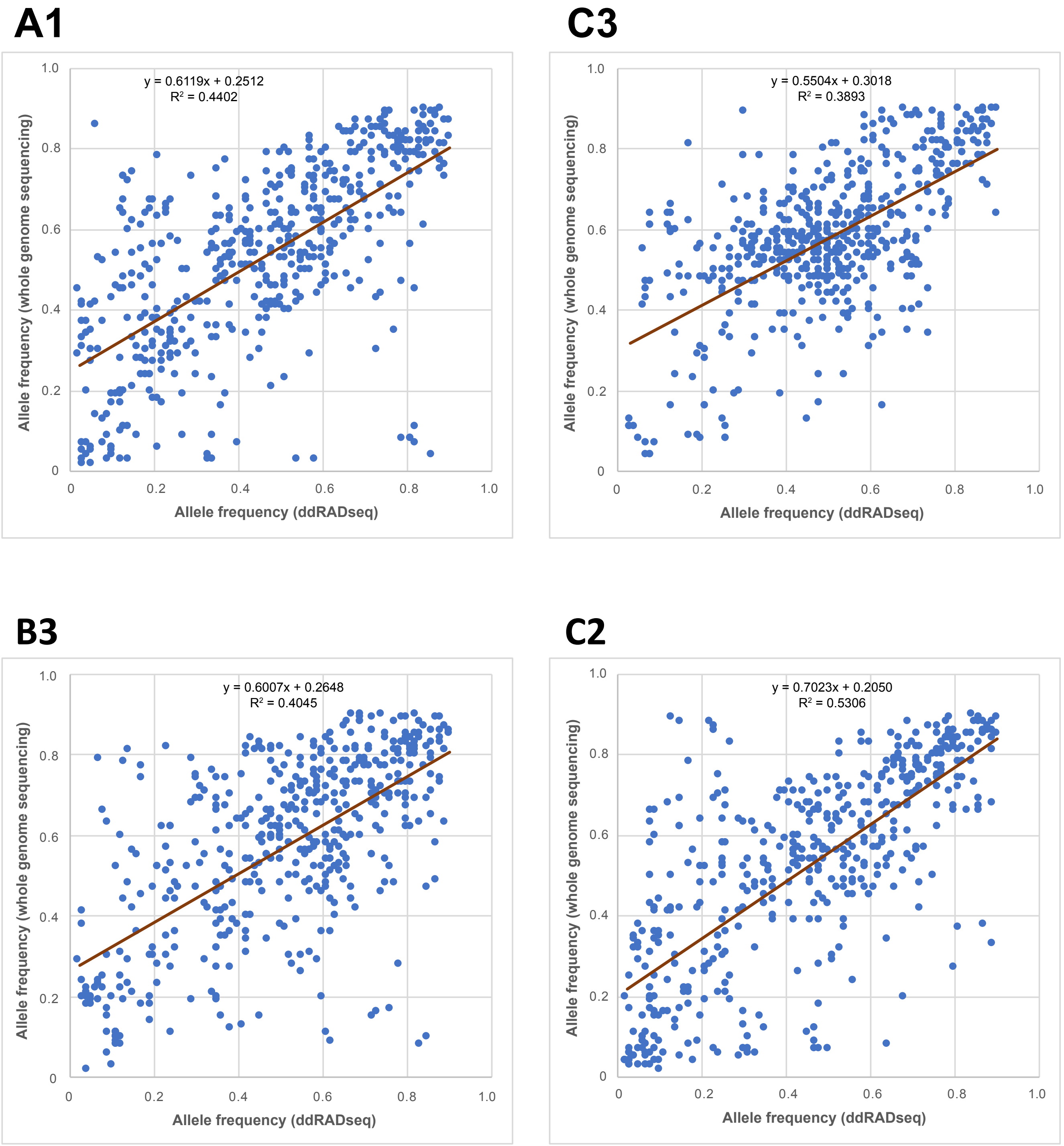


**Figure S8.** Scatterplots showing the relationship between allele frequency at bi-allelic sites observed in whole genome sequencing data and ddRADseq data from five isolates of *R. irregularis* (A1, C3, A5, B3 & C2).

**Figure S9. Analysis of frequencies at bi-allelic sites in isolate DAOM197198.**

**a** Allele frequency plots when assembly DNA1 was used as a reference genome
(only coding and non repeated regions)

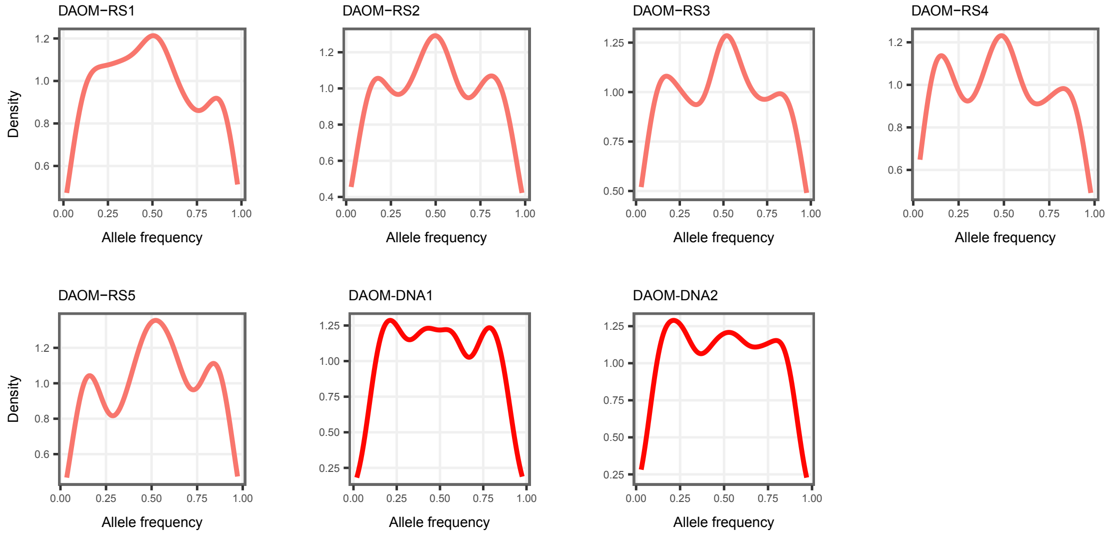


**b** Allele frequency plots when assembly DNA2 was used as a reference genome
(only coding and non repeated regions)

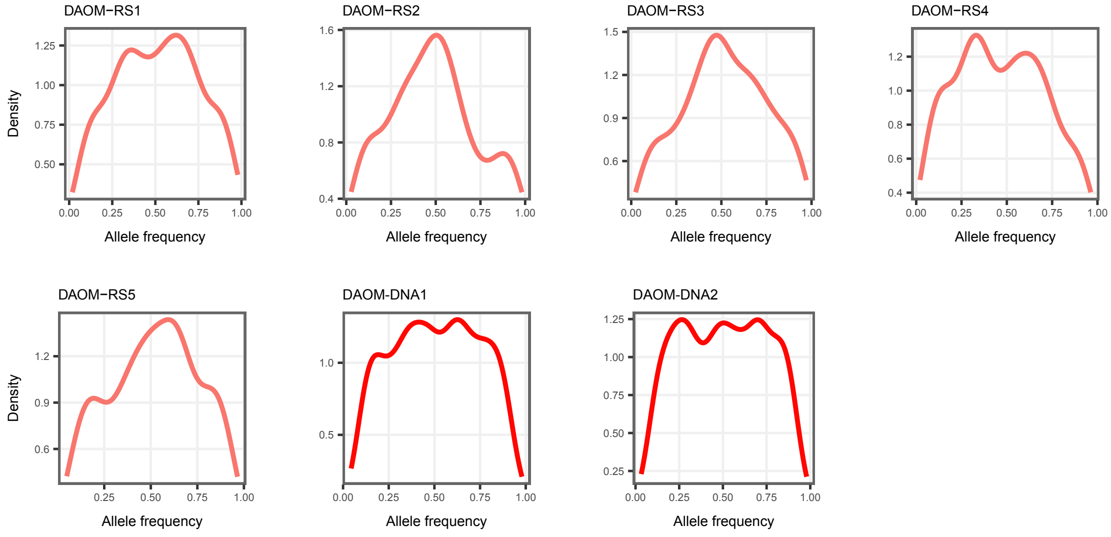

**c** Boxplot of depth of coverage for ddRAD-seq data and WG data of the isolate DAOM197198.

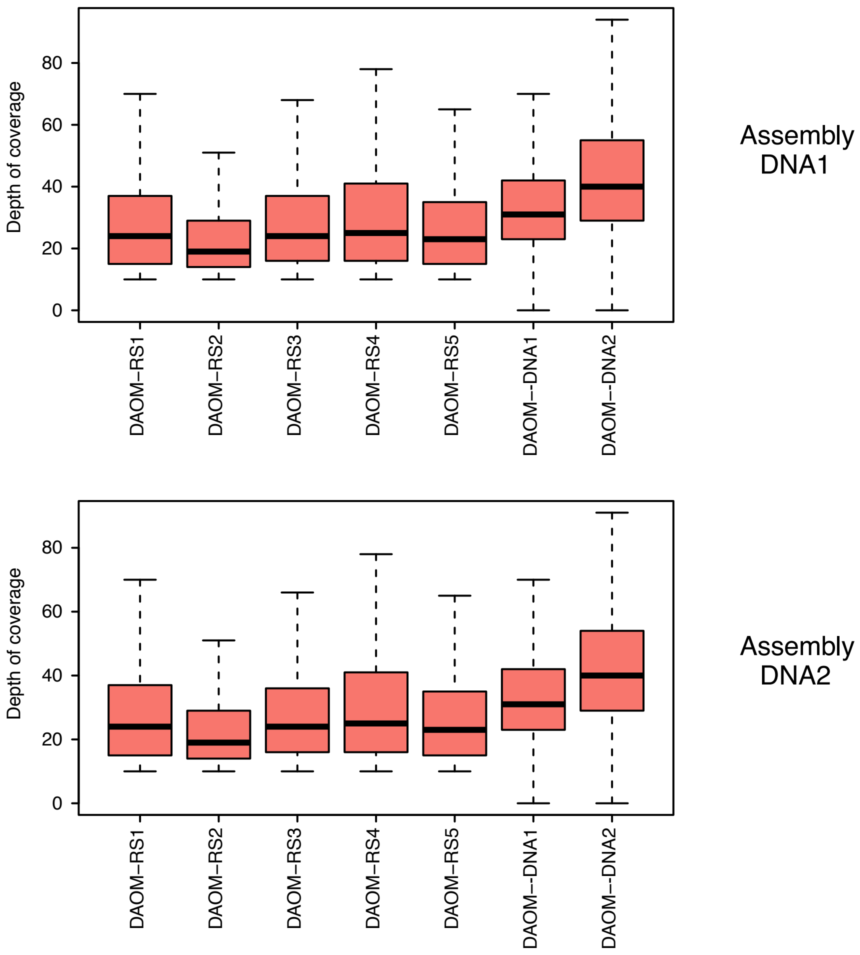


**d** Mean number of bi-allelic positions per 1 kb region versus depth of coverage detected in WG data in non-repeated and non-coding regions. This figure is the same as figure 2b with the addition of DAOM197198 ddRAD-seq and WG samples (pink).

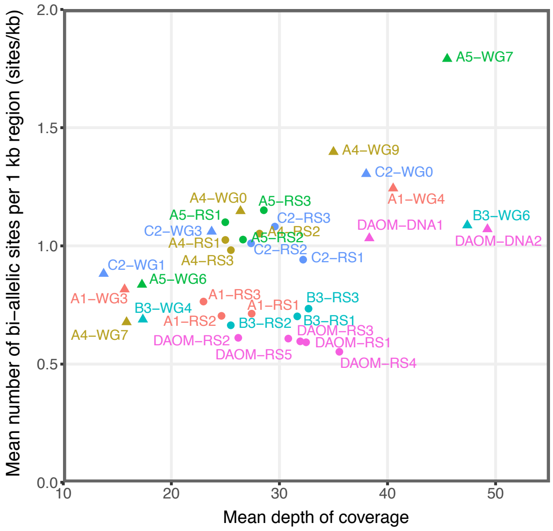

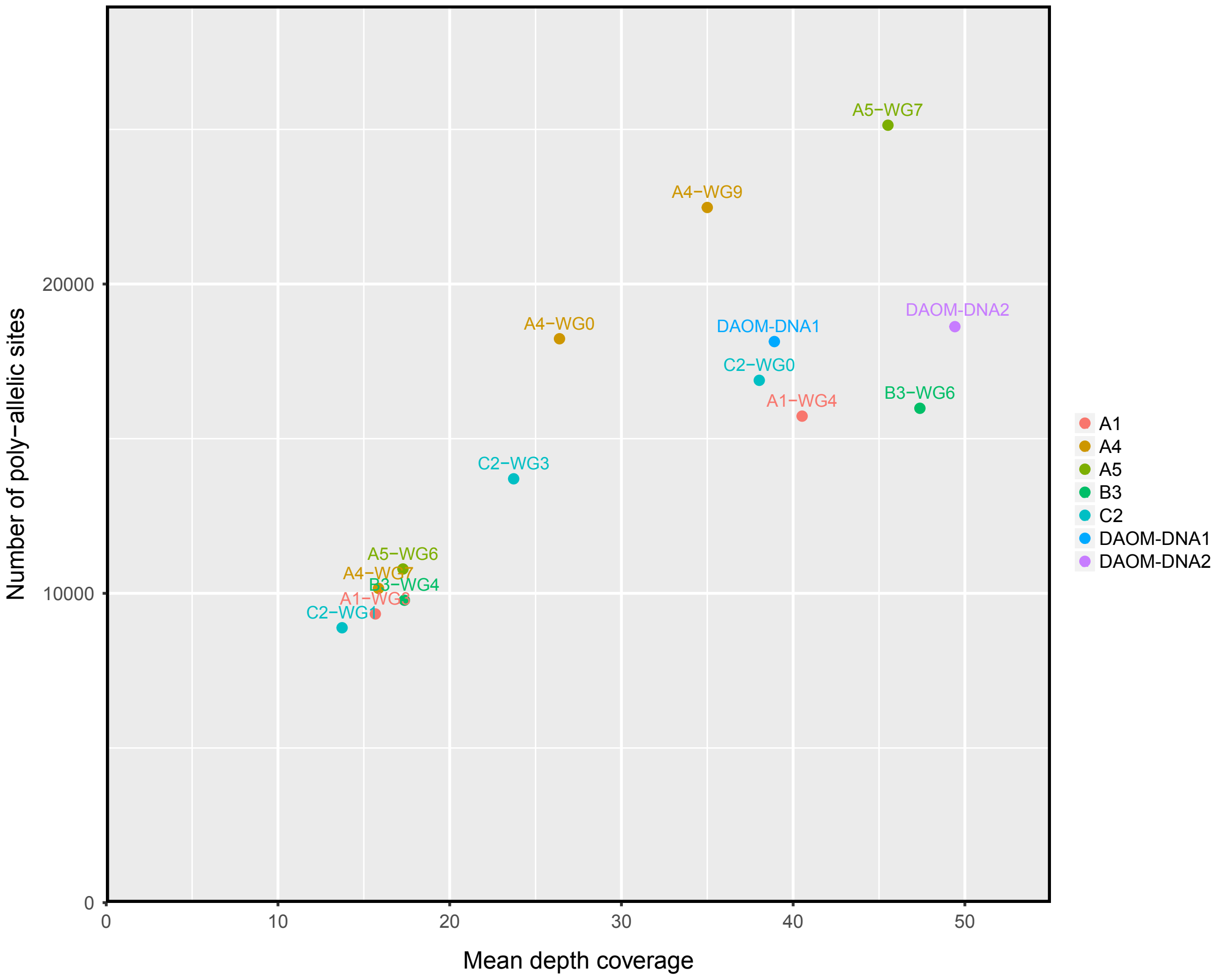


**Figure S10: Mean number of bi-allelic positions versus mean depth of coverage detected in whole genome (WG) data in non-repeated and coding regions across the whole genome assembly.** The whole genome assembly was considered and not only the predicted ddRAD-seq regions.

**Figure S11: Allelic frequency graphs obtained with different depth coverage thresholds.**
All positions in the genome, except the regions labelled as 'repeats', were considered for this analysis. Randomly chosen frequencies of one allele at di-allelic positions were considered in order to generate the distribution. Only positions with a total depth coverage higher or equal to the threshold were included.

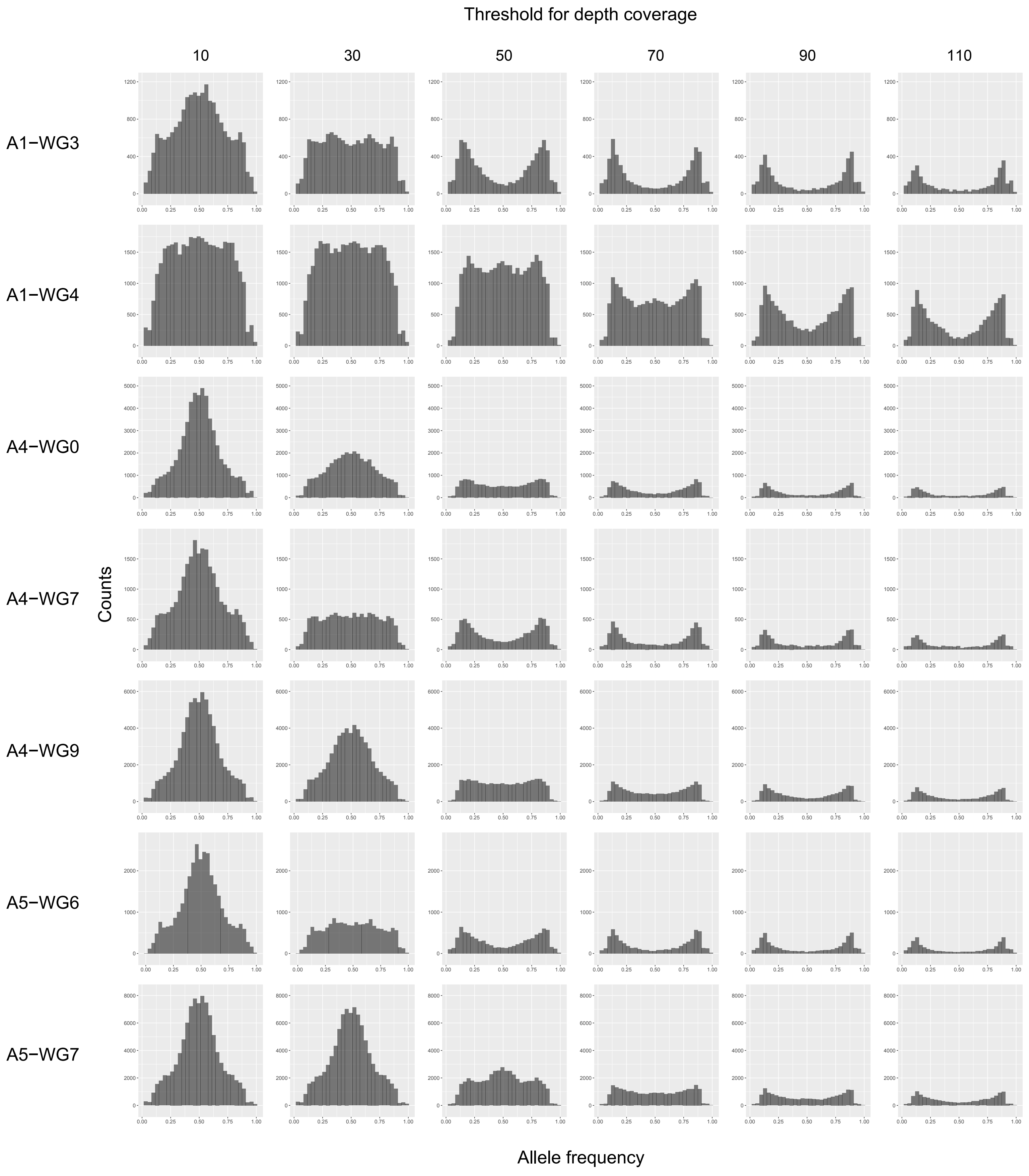

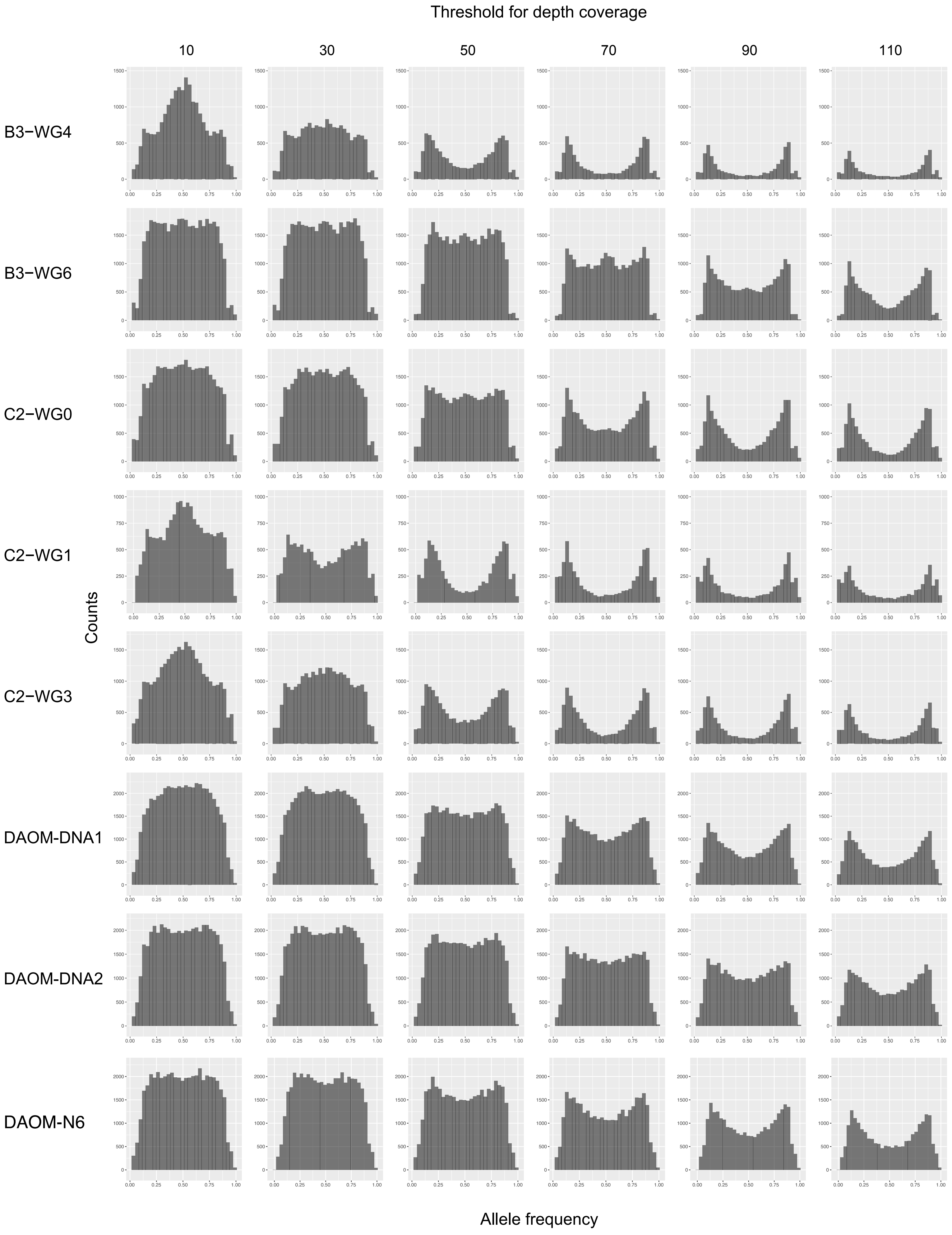

Threshold for depth coverage
10
30
50
70
90
110
B3−WG4
B3−WG6
C2−WG0
C2−WG1
C2−WG3
DAOM-DNA1
DAOM-DNA2
DAOM-N6
Counts
Allele frequency

**Figure S12: Analysis of RNAseq data in other fungi and plants**

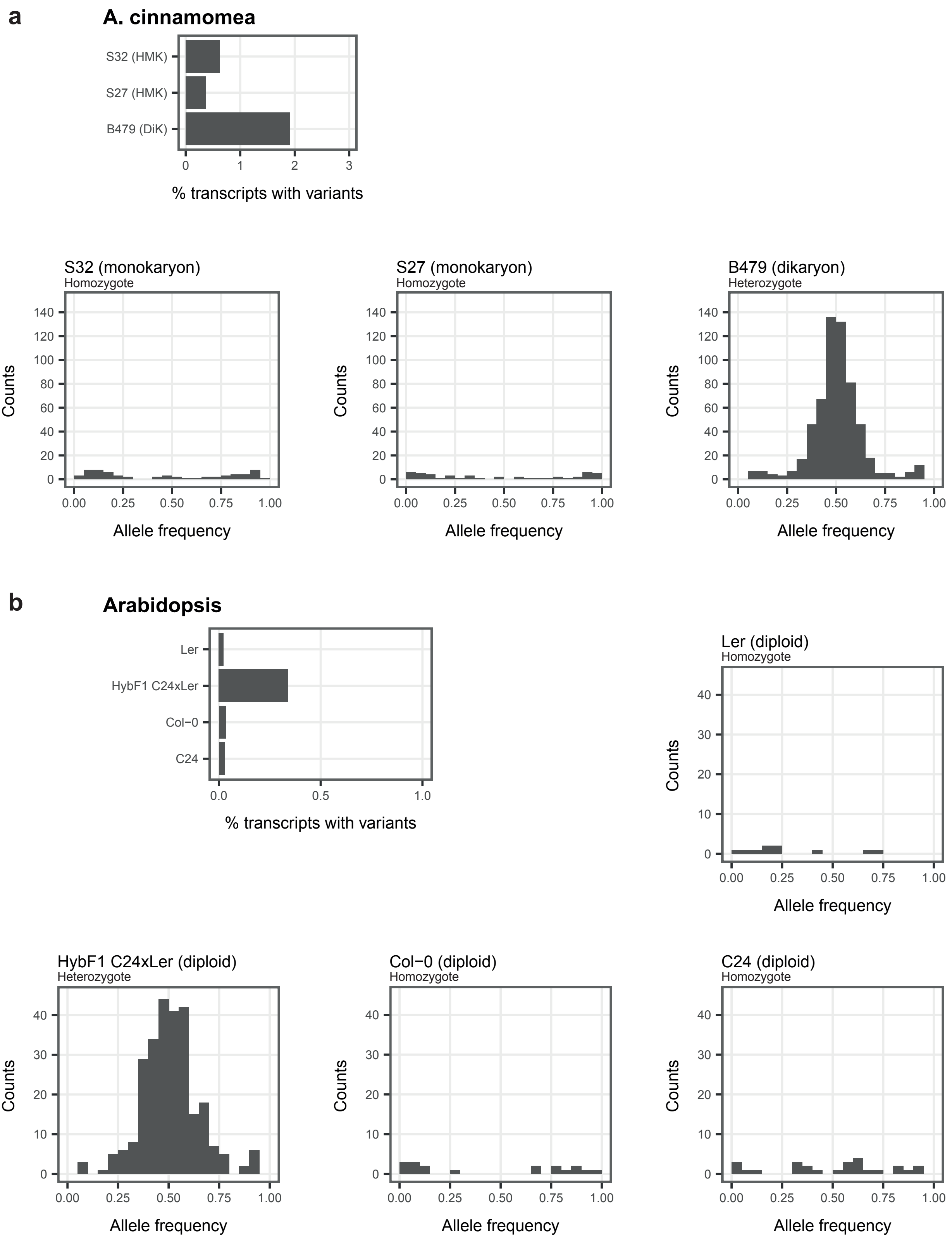

**c** **Plants (Wheat and precursor species)**

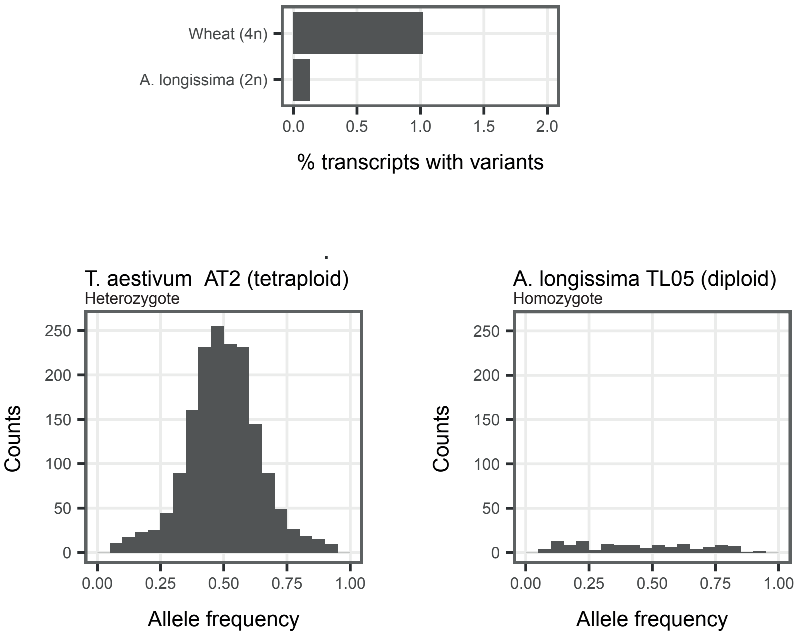


**d** **Unicellular fungal species**

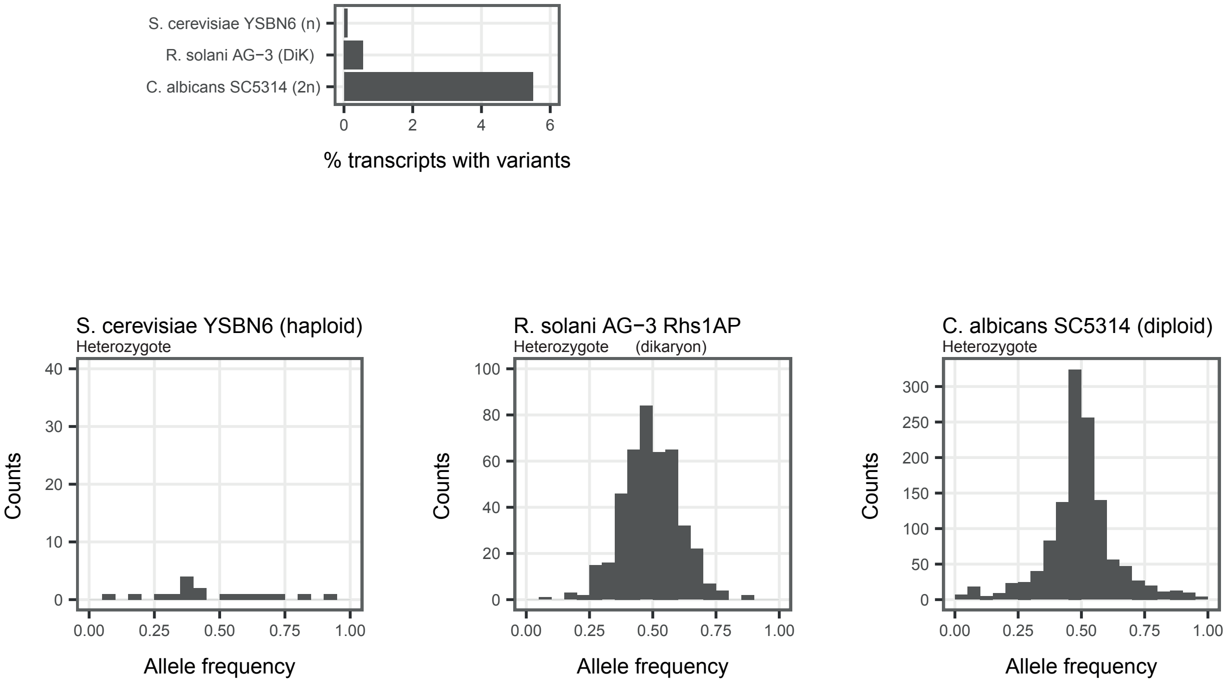

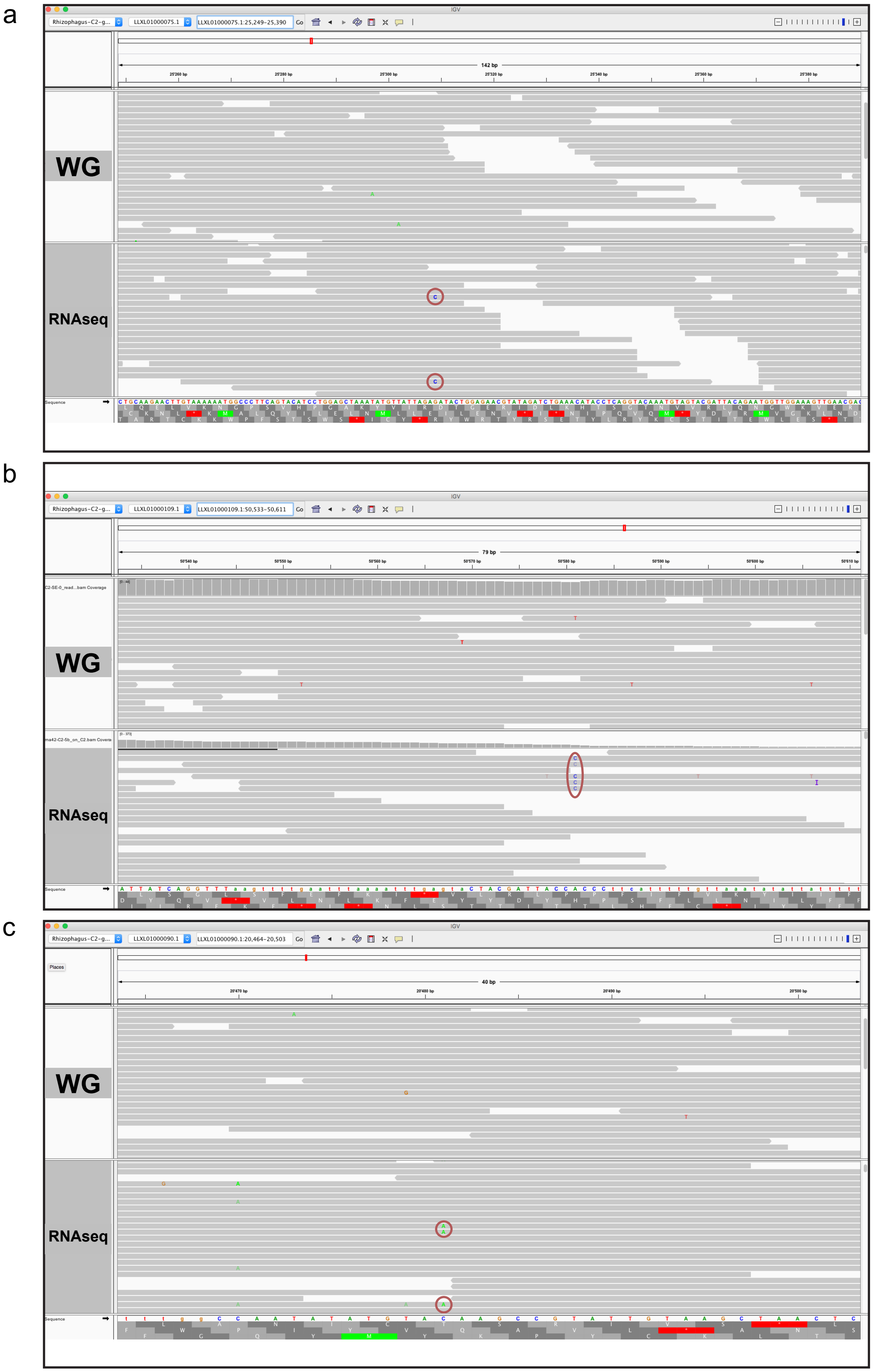


**Figure S13: Examples of bi-allelic positions found in RNA-seq data and not in WG data in isolate C2.** (a), (b) and (c) correspond to three examples in different regions of the genome displayed in IGV browser. Only nucleotides differing from the reference genome are shown with a colour different from light gray. Alleles contributing to bi-allelic positions in RNA-seq are surrounded with red circles or ellipses.

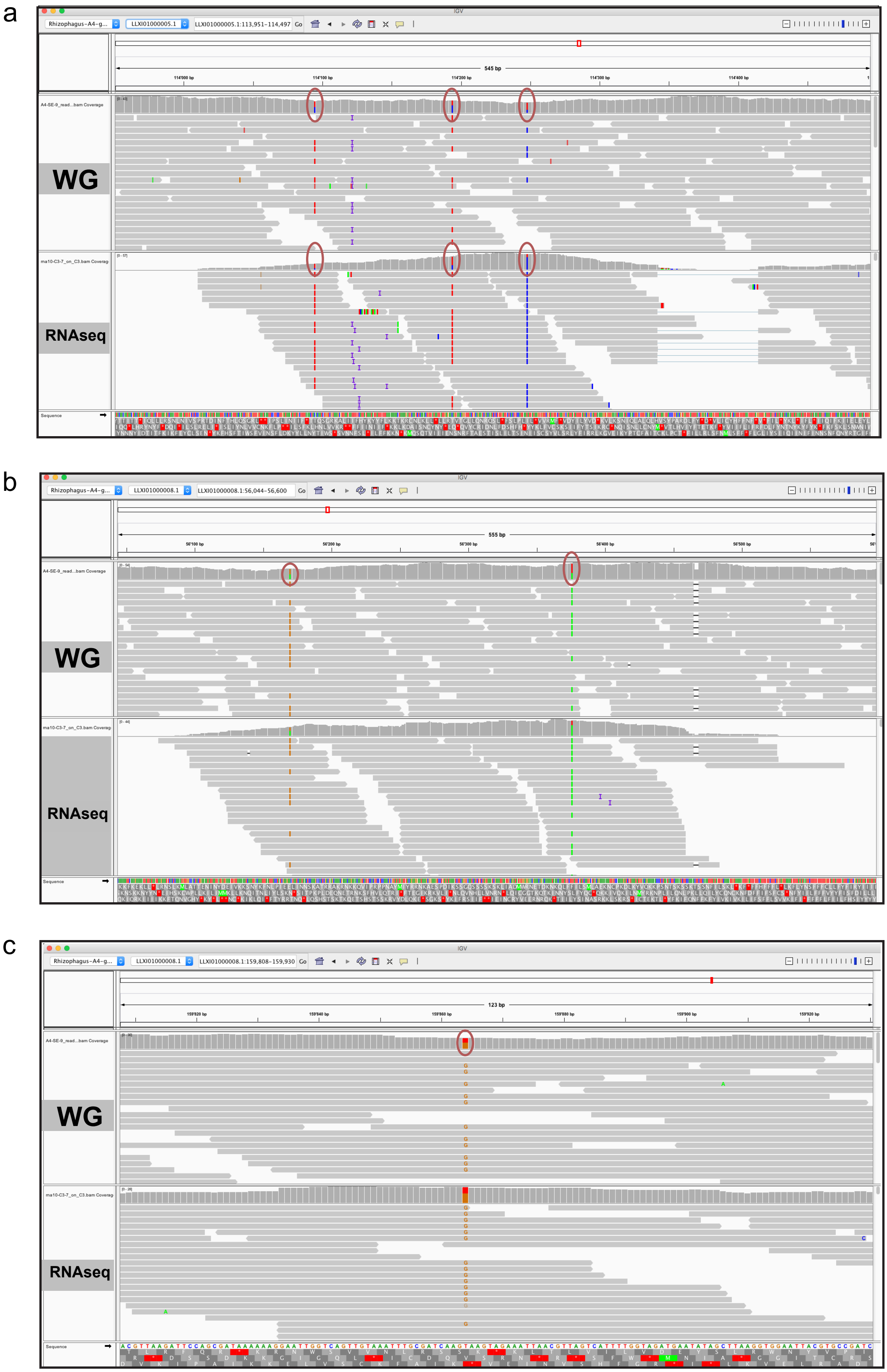


**Figure S14: Examples of bi-allelic positions found both in RNA-seq data and in WG data in isolate C3.** (a), (b) and (c) correspond to three examples in different regions of the genome displayed in IGV browser. Only nucleotides differing from the reference genome are shown with a colour different from light gray. Alleles contributing to bi-allelic positions in RNA-seq are surrounded with red circles or ellipses.

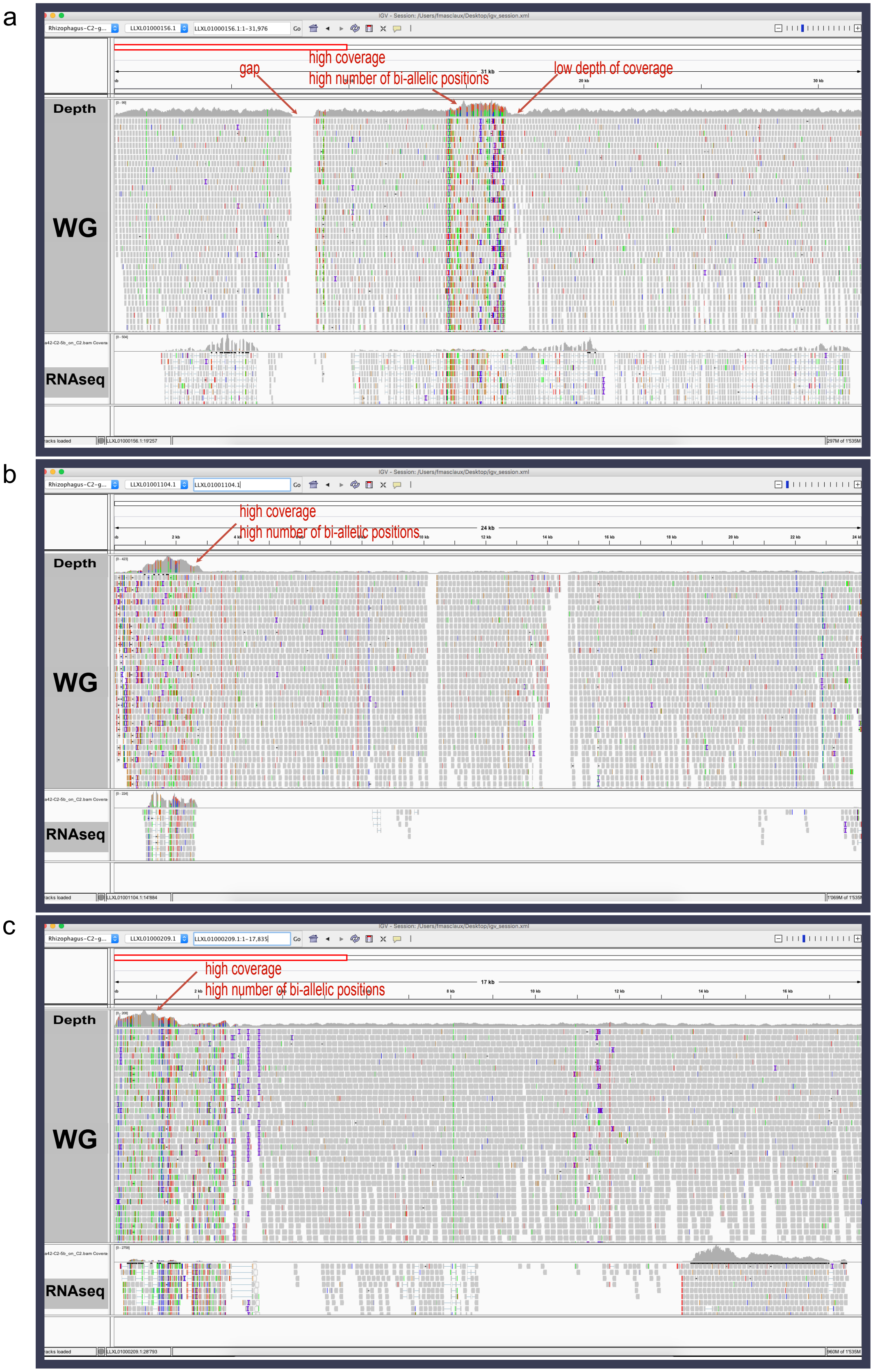


**Figure S15: Examples of problematic regions in the genome assembly of the isolate C2.** (a), (b) and (c) correspond to three examples in different regions of the genome displayed in IGV browser. Colours other than light grey depict alleles different from the reference in the reads. 'Depth' shows a histogram of the depth of coverage for genomic reads. Interesting regions are labelled with red text and boxes.
